# Assessing the burden of rare DNA methylation deviations in schizophrenia

**DOI:** 10.1101/2023.01.09.523073

**Authors:** Christine Søholm Hansen, Andrew McQuillin, David St Claire, Jonathan Mill, Eilis Hannon, Andrew J. Sharp, Magdalena Janecka

**Author notes:** Corresponding authors: Christine Søholm Hansen, PhD, Department of Psychiatry, Icahn School of Medicine at Mount Sinai, Magdalena Janecka, PhD, Department of Psychiatry, Icahn School of Medicine at Mount Sinai.

## Abstract

Along with case-control group differences in DNA methylation (DNAm) identified in epigenomewide association studies (EWAS), multiple rare DNAm outliers may exist in subsets of cases, underlying the etiological heterogeneity of some disorders. This creates an impetus for novel approaches focused on detecting rare/private outliers in the individual methylomes. Here, we present a novel, data-driven method - Outlier Methylation Analysis (OMA) – which through optimization detects genomic regions with strongly deviating DNAm levels, which we call outlier methylation regions (OMRs).

Focusing on schizophrenia (SCZ) - a neuropsychiatric disorder with a heterogeneous etiology – we applied the OMA method in two independent, publicly available SCZ case-control samples with DNAm array information. We found SCZ cases had an increased burden of OMRs compared to controls (IRR=1.22, p=1.8×10^-8^), and case OMRs were enriched in regions relevant to cellular differentiation and development (i.e. polycomb repressed elements in the Gm12878 differentiated cell line, p=1.9×10^-5^, and poised promoters in the H1hesc stem cell line, p=5.4×10^-4^). Furthermore, SCZ cases were ~2.5-fold enriched (p=1.1×10^-3^) for OMRs overlapping genesets associated with developmental processes. The OMR burden was reduced in clozapine-treated, compared to untreated, SCZ cases (IRR=0.88, p=9.5×10^-3^), and also associated with increased chronological age (IRR=1.01, p= 2.7×10^-16^).

Our findings demonstrate an elevated burden of OMRs in SCZ, implying methylomic dysregulation in SCZ which could correspond to the etiological heterogeneity among cases. These results remain to be causally examined and replicated in other cohorts and tissues. For this, and applications in other traits, we offer the OMA method to the scientific community.

## Introduction

Schizophrenia (SCZ) is a neuropsychiatric disorder with a lifetime prevalence of 0.7% in the general population ^1^, which can have considerable impact on a person’s thoughts, perception and behavior. It is diagnosed based on clinically observable or reported symptoms, such as delusions, hallucinations, apathy, and lack of pleasure, but is often associated with heterogeneous clinical manifestations, ages of onset and familial histories ^2^.

Twin studies indicate that around 80% of phenotypic variance of SCZ is attributable to genetic factors^3^. These factors include variants across the full allele frequency spectrum, including both common^4^ and rare variants (usually found in < 1% of the population). Previous studies have demonstrated an increased burden of rare coding variants in loss-of-function intolerant genes, and ultra-rare structural variants in SCZ cases compared to controls^5–8^. Cases with such rare, high-impact variants have on average lower polygenic risk score – underlain by common disease variants – than cases without such variants^9^. These observations suggest that SCZ may have diverse genetic etiologies, spanning a polygenic burden of many common variants, as well as high-impact, rare variants. Additionally, environmental factors relating to prenatal exposures, childhood adversities or lifestyle are associated with SCZ and may interact with the genetic risk^10^.

Both genetic and environmental influences can impact the epigenome – molecular changes ‘on top’ of the genome (epi ~ *above*). These can, along with other functions, regulate gene expression and orchestrate cellular differentiation and development. A widely studied epigenetic modification is DNA methylation (DNAm), which is vital for genomic stability and gene silencing^11^ as well as embryonic development and response to environmental stimuli^12^. The evidence for the involvement of the methylome in SCZ pathology, comes from several sources, including (i) the co-occurrence of common SCZ genetic risk variants in sites impacting DNAm (i.e. methylation quantitative trait loci (mQTLs)) in the fetal brain^13^, and (ii) the identification of differentially methylated positions (DMPs) between SCZ cases and controls ^14^ in epigenomewide association studies (EWAS) – some of which may arise as a consequence, rather than cause of SCZ.

Complementary to studies focused on common DNAm variation in SCZ (i.e. EWAS), is the analysis of rare epigenetic events found only in small subsets of cases. Rare outlier DNAm events may, similarly to rare *genetic* variation, underlie etiological heterogeneity of SCZ, offering insights into the disease etiology and even explaining cases with no discernible genetic cause (i.e. stochastic or environmental impacts). Additionally, their association with rare genetic variants^15^ could offer insights into the disease mechanisms in carriers of such rare variants.

Previous studies have detected very deviant and ultra-rare methylomic variation (referred to as *epivariations*) by querying the individual methylomes^16^. Such *epivariations* have been linked with substantial changes in gene expression^16^ and found to be enriched in disease-relevant genes in SCZ cases^17^. However, the role of rare, but moderate deviations of DNAm levels, remains unknown, and calls for a new approaches to detect and quantify them. To address this, we have developed a novel method and used it to systematically detect and quantify outlier methylation across a wider range of deviation (from modest to extreme deviation levels) in two independent SCZ case-control samples.

Our method, *Outlier Methylation Analysis* (OMA), uses optimization to systematically detect regions with similar methylomic deviation from the cohort median – which we refer to as outlier methylation regions (OMRs). In the optimization we leveraged the co-methylated nature of the proximal CpG sites^18^ to detect regional DNAm deviations. Our method allows to robustly detect OMRs, analyse their burden related to the case-control status, and stratify them by their overall deviation, genomic size and regional context - even if such events are only found only in a single individual.

Similar to testing the composite burden of rare genetic variants in SCZ cases, our aim was to assess the individual OMR burden in SCZ cases and controls - both from their overall number, and stratified by their properties (i.e. direction, size and deviation). We further assessed specific functional enrichments of OMRs in SCZ cases compared to controls, and performed gene-set overrepresentation analysis of OMRs overlapping genes in the individual methylomes.

## Methods

### Data

We used two publicly available SCZ case-control datasets: the University College London (UCL)^19^ [GEO accession: GSE80417] consisting of 322 controls and 353 SCZ cases, and the Aberdeen data^20^ [GEO accession: GSE84727], consisting of 433 controls and 414 SCZ cases (for detailed description of both see^14^). These datasets include demographic (age and sex) and clinical (SCZ diagnosis and clozapine treatment) information, and genome-wide DNA methylation (DNAm) data profiled from whole blood using Illumina Infinium 450k Human Methylation beadchip v1.0 (450k). The treatment (clozapine, yes/no) information was only available for a subset of SCZ cases in each dataset (UCL: 32.0% and Aberdeen: 70.0% of SCZ cases). For each cohort, we calculated DNAm-based estimates of blood cell proportions (DNAmBCPs) for 5 major blood cell types with the reference panel described in ^21^ (excluding NK cells, which in most samples yielded a proportion of 0) using the ENmix R-package ^22^ and DNAm-based smoking scores (DNAmSS) with the smoking EWAS summary statistics from ^23^ and the method described in ^14^. We tested case-control differences in demographic and methylome-derived variables with a Welch 2 sample T-test for continuous data or Fisher’s exact test for binary data. All analyses were performed using R software ^24^.

### QC and filtering

The methylomic data quality control (QC) of each cohort is described in ^14^. In short, samples were excluded if they had mean methylated or unmethylated intensities < 2500, or median bisulfite conversion controls probes < 90, were a mismatch between estimated and reported sex, were principal component (PC) outliers (PC1 or PC2 > 2 SDs from the mean PC1 and PC2) or had detection p-values (detP) > 0.05 across more than 1% of probes (i.e. call rate < 99%). We further removed slides containing fewer than 2 samples or only cases or controls (5 samples in the UCL cohort and 6 samples in the Aberdeen cohort were removed) and excluded probes if they had detP > 0.05 in more than 1% of samples (i.e. call frequency < 99%), were non-CG or X/Y chromosomal probes, or were identified as cross hybridizing probes in ^25 26^ with the *maxprobes* R-package for the 450k array.

### Data preprocessing

For each dataset we normalized the filtered methylated and unmethylated intensity data with the *dasen* method in the *wateRmelon* R-package ^27^, then calculated the DNAm proportions (beta) values and logit transformed these into M-values (see Supplementary Methods Figure 1A-B). We then linearly adjusted the M-values for variance associated with the 5 DNAmBCPs described above and the first 10 PCs. Finally, we performed an outlier robust Z-score transformation by centering the M-values with their median and scaling them with their median absolute deviation (MAD) * 1.4826 at each site. The derived values approximate standard deviations (SD) from the cohort median at each site (see Supplementary Methods Figure 1B-C).

The method for OMR detection - *Outlier Methylation Analysis* (OMA) - is described in more detail in the Supplementary Methods. In short, we query the individual methylome for regions with similar levels of DNAm deviation from the norm (Z-scores), by applying an optimization of Stouffer-Liptak combined Z-scores (CombZ), described in ^28^. To reduce computational demands, we conditioned the detection on sites deviating by at least +/− 3 SDs and separated by no more than 1000 bps and no more than one “non-outlier” site (i.e. sites with < 3 and > −3 SD deviation). Finally, we removed OMRs with majority of the underlying CpGs outlying in more than 5% of samples.

### Association between the OMR burden and clinical phenotypes

We analyzed SCZ casecontrol differences in (**a)** total OMR burden, and in OMR burden stratified by (**b)** direction (positive and negative, corresponding to hyper- and hypomethylated regions, respectively), (**c)** deviation from the norm (i.e OMR absolute mean Z-score (|MeanZ|) deviation bins, ranging from 3 SDs to >8 SDs in increments of 1 SD) and (**d)** genomic size (from 0bp up to >800bp, in increments of 100bp). While (a) allows us to consider differences in the overall OMR burden, in (b) – (d) we consider if case-control differences were associated with specific OMR properties.

We tested over-dispersion of the count data with the *dispersiontest* function in the AER R-package ^29^ and used a negative binomial (NB) model (for over-dispersed count data - where the variance is significantly larger than the mean), to test case-control differences in OMR burden. If the NB model failed to converge in 50 iterations, we used a simpler Poisson model. We used the glm.nb function in *MASS* R-package ^30^ for the NB model and the *glm* function in the *stats* R-package for the Poisson model.

We detected samples with extreme burden of OMRs using the *aout.nbinom* function in the *alphaOutlier* R-package ^31^, and excluded them from each analyses. For this alpha outlier detection we applied the default alpha=0.1 cutoff to the data distribution, fitted with a negative binomial model with *fitdistr* function in the MASS R-package ^30^. While these samples were not included in the burden analyses to preclude their undue influence on the results, to assess biological relevance of the alpha outlier samples we tested their enrichment of SCZ cases using the Fisher’s exact test.

All models were run first without adjustment (crude), and then iteratively adjusted for the covariates: **i)** DNAmSS+age+sex, and **ii)** DNAmSS+age+sex+clozapine. As adjustment resulted in sample attrition due to missing covariate information, we repeated the crude analysis in a reduced sample having the full covariate data, to assess if parameters’ change was due to sample attrition or covariate adjustment. Since global confounders of DNAm levels (batch effects and cellular composition in the bulk blood tissue) had already been linearly adjusted for, we did not further adjust for them here.

We obtained OMR incidence rate ratios (IRR) – interpretable as change in OMR burden associated with case/control status – by exponentiating the NB model estimates and obtaining their corresponding 95% confidence intervals. We adjusted the p-values for multiple testing using the false discovery rate (FDR) correction within each analysis type a)-d) across all iterative adjustments i)-iii) and included covariates, using the *p.adjust* function in the *stats* R-package. Statistical significance was set at FDR < 0.05.

### OMR (epi)genomic feature enrichment

We annotated OMRs to genomic and epigenomic features with annotation tracks available in the R-package *annotatr ^32^* for the hg19 genome build, i.e. CpG islands (CGIs) (*islands, shores, shelves* and *inter-CGI*), genic features (*promoters, exons/introns, 5UTRs, 3UTRs, 1to5kb* and *intergenic*) and active chromatin states (ChromHMM) (*ActivePromoter, WeakPromoter, PoisedPromoter, StrongEnhancer, WeakEnhancer*, *Insulator*, *TxnElongation*, *TxnTransition*, *WeakTxn*, *Heterochrom/lo*, *Repressed* and *Repetitive/CNV*) measured in nine cell lines ^33^. For the genic annotations, we combined *introns* and *exons* into one feature (*exons/introns*), and only annotated OMRs to genic features of genes that had a gene entrez id, and otherwise as *intergenic*. Each OMR is annotated with at least one feature within each track. To assess enrichment or depletion of OMRs in specific (epi)genomic features, we used a linear model to analyze the proportion of an individual’s OMRs annotated to each feature. We adjusted for covariates found to associate with the OMR burden in the previous analysis, and only stratified OMRs by direction (positive and negative). P-values were FDR-adjusted for multiple testing within the genomic (CGI and genic) and epigenomic (all nine ChromHMM) annotation tracks, across all iterative adjustments i)-iii) and covariates. Statistical significance was set at FDR<0.05.

### Gene ontology overrepresentation analysis

We performed gene set overrepresentation analysis (ORA) of OMRs overlapping genes, independently in each individual. First, we used the *gsaregion* function from the *missmethyl* R-package to assign OMRs to genes. The *gsaregion* function first reduces OMRs to individual sites and assigns them to gene(s) – while adjusting for bias relating to gene probe number density and multiple gene probes from overlapping genes – and then performs hypergeometric testing for gene set overrepresentation ^34^. We used the GO biological pathway gene sets from MSigDB, and aggregated all *significant* GO-terms into their 21 hierarchical parent terms (*goslim_agr* on geneontology.org) defined with the *goSlim* function in the *GSEABase* R-package. Some terms had no parent term and were instead included in a miscellaneous (mics) category. We used logistic regression to test for enrichment of SCZ cases in the 21 GO-terms, misc and any category, while adjusting for the covariates significantly associated with OMR burden). To avoid sparse data bias, we only included GO terms significant in at least 10 individuals. P-values were FDR-adjusted for multiple testing. Statistical significance was set at FDR<0.05.

### Meta-analysis

We used fixed effect models implemented *meta* R-package ^35^ to meta-analyze the model estimates and standard errors derived from both cohorts. We applied FDR multiple testing correction within each analysis and set statistical significance at FDR < 0.05. We excluded the meta-analysis results with evidence for significant heterogeneity between the samples (heterogeneity p-value (hetP) < 0.05).

## Results

### Sample characteristics

After QC the DNAm datasets contained 432,130 sites for 670 samples in the UCL cohort (in 319 controls and 351 cases), and 432,366 sites for 841 samples in the Aberdeen cohort (in 433 controls and 408 cases). The UCL controls included fewer males and were on average younger than cases (Table 1). In both cohorts SCZ cases had significantly higher DNAm smoking scores (**DNAmSS**) and differed from controls in their DNAm-based estimates of blood cell proportions (**DNAmBCP**) (Table 1; these differences have been reported in ^36^).

**Table 1:** Samples’ demographics and DNA-derived parameters. We tested differences between cases and controls using a T-test (continous variable) or Fisher Exact Test (binary). Statistically significant differences are highlighted with bold for p<0.05. P-values: (*)<0.1, *<0.05, **<0.01, ***<0.001, ****<0.0001. DNAmSS – DNAm-derived smoking score; CD4T/CD8T – estimated fraction of CD4/CD8 lymphocytes; Gran - estimated fraction of granulocytes; Mono - estimated fraction of monocytes; all cell fractions were estimated in blood.

|  | UCL cohort |  |  | Aberdeen cohort |  |  |
| --- | --- | --- | --- | --- | --- | --- |
|  | SCZ<br>(N=351) | Control<br>(N=319) | P-value | SCZ<br>(N=408) | Control<br>(N=433) | P-value |
| age (mean+/-SD) | 43.61+/-14.66 | 36.84+/-14.64 | 1.09E-08**** | 44.31+/-13.95 | 44.87+/-12.16 | 5.99E-01 |
| Sex (M%) | 0.72 | 0.44 | 3.84E-13**** | 0.68 | 0.74 | 8.08E-02(*) |
| Clozapine (Yes%) | 0.49 | 0 | NA | 0.33 | 0 | NA |
| DNAmSS (mean+/-SD) | 57.46+/-1.45 | 56.36+/-1.21 | 7.06E-25**** | 49.41+/-2.15 | 47.97+/-1.74 | 1.24E-24**** |
| Bcell (mean+/-SD) | 0.07+/-0.03 | 0.07+/-0.02 | 8.43E-01 | 0.06+/-0.03 | 0.06+/-0.02 | 7.38E-01 |
| CD4T (mean+/-SD) | 0.14+/-0.07 | 0.14+/-0.05 | 6.84E-01 | 0.12+/-0.06 | 0.13+/-0.05 | 3.18E-06**** |
| CD8T (mean+/-SD) | 0.1+/-0.05 | 0.13+/-0.05 | 8.89E-18**** | 0.1+/-0.05 | 0.13+/-0.05 | 2.52E-17**** |
| Gran (mean+/-SD) | 0.65+/-0.09 | 0.61+/-0.07 | 9.68E-10**** | 0.66+/-0.08 | 0.61+/-0.07 | 1.15E-20**** |
| Mono (mean+/-SD) | 0.06+/-0.03 | 0.07+/-0.02 | 6.79E-01 | 0.07+/-0.02 | 0.06+/-0.02 | 1.88E-01 |

### OMR burden distribution

We detected a total of 93,553 OMRs in the UCL cohort and 105,746 OMRs in the Aberdeen cohort with absolute overall deviations (|MeanZ|) between 3-27.9 SDs and 3-31.9 SDs, respectively (see Supplementary Figures S2-7). The median overall OMR burden per sample was 82 in both cohorts, ranging between 24-1,598 and 32-1,498, respectively, and their distribution was over-dispersed (overdispersion = 202.8, p=3.6×10^-7^ and 169.6, p=4.3×10^-7^, respectively), validating the use of a negative binomial (NB) model to fit the data.

Some samples had an abnormally large OMR burden - as indicated by an alpha outlier cutoff of 0.1 (see methods). We detected 67 and 66 alpha outlier samples defined at >292 and >246 OMRs/sample, respectively (Supplementary Figure S8). These samples were enriched for SCZ cases in both cohorts (Fisher OR = 1.6, p = 0.09 and 2.3, p=0.003, respectively). Since NB estimates may be sensitive towards the inclusion of extreme outlier values, we removed these from the main analyses, but performed sensitivity analyses where we retained them.

### The burden of OMRs is increased in SCZ cases, but reduced in association with clozapine treatment

We found a significantly higher OMR burden in SCZ cases compared to controls in both cohorts (IRR [95%CI] = 1.30 [1.19:1.41], p=5.24×10^-9^ and 1.17 [1.11:1.24], p=5.20×10^-8^) and in the meta-analysis (Table 2; Supplementary Tables, Meta Tab 2). These results remained significant after covariate adjustment (Figure 1A and D; Supplementary Tables GSE80417 and GSE84727 Tab 2) and retaining the alpha outlier samples (Supplementary Figure S9). Heterogeneity between the two cohorts was not statistically significant (Table 2).

**Table 2:**
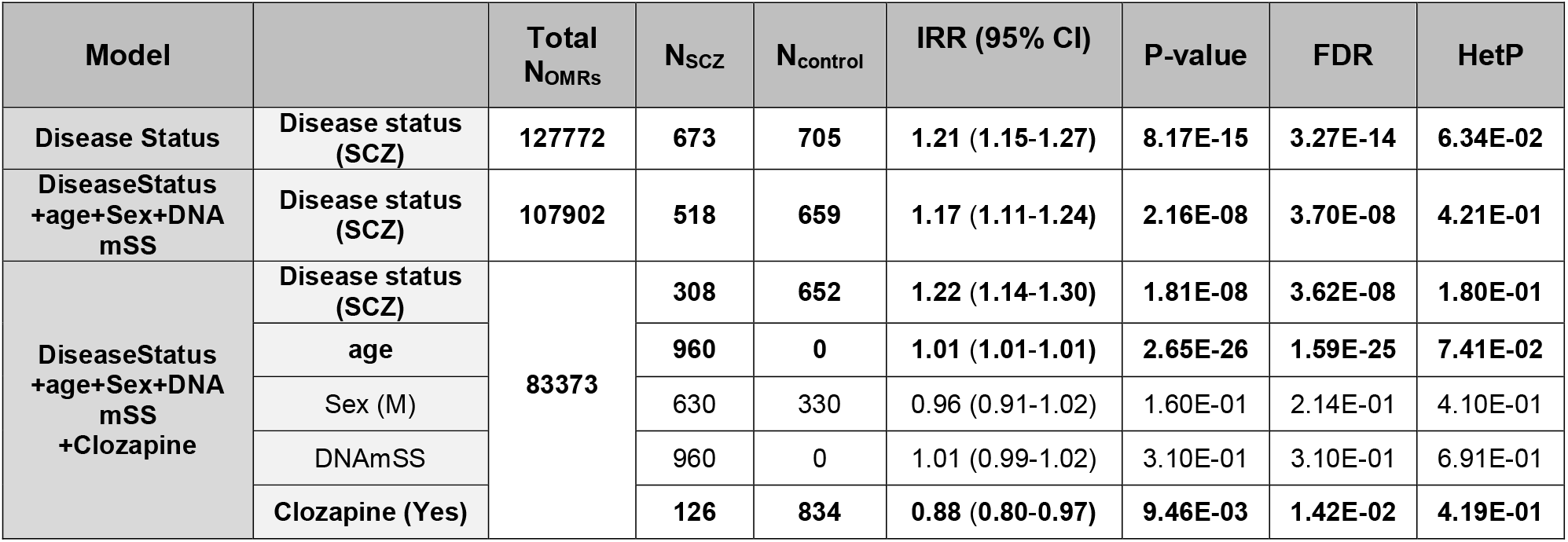
Fixed effects meta-analysis of the crude and adjusted associations between schizophrenia (SCZ) and the total OMR burden in the two cohorts. Covariates included in the adjusted models were: chronological age, sex, DNAm based smoking scores (DNAmSS) and clozapine use. Outlier samples have been removed and the number of samples included is indicated as N_scz_ and N_control_. Effect size estimate is given as the incidence rate ratio (IRR) with the 95% CI [2.5%:97.5%], and the heterogeneity p-values (HetP) indicate significant difference between the two cohorts. P-values were adjusted for multiple testing with FDR. Variables significantly associated with OMR burden (FDR<0.05) are indicated with bold.

**Figure 1:**
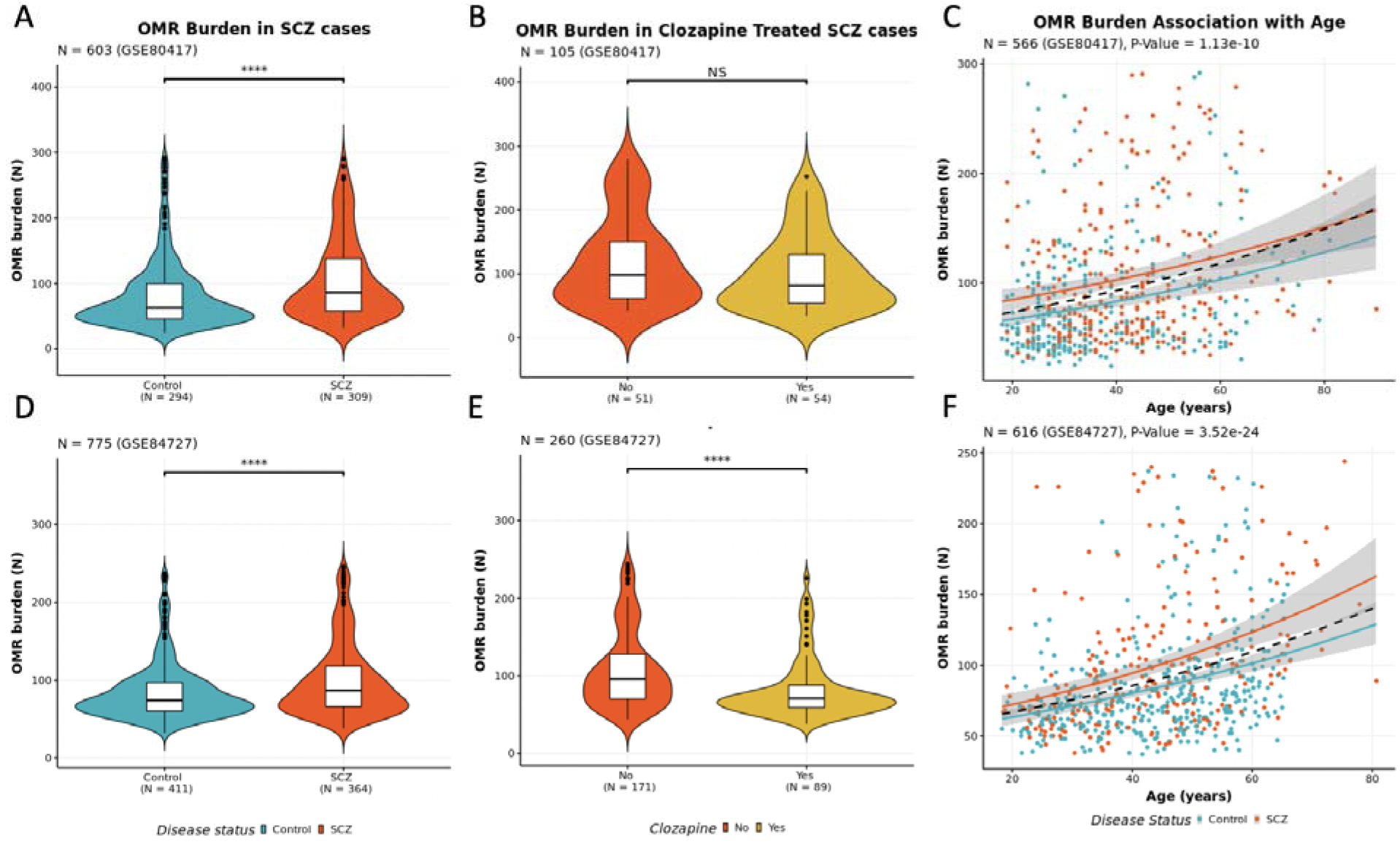
OMR burden association with SCZ (A and D), clozapine treatment (D and E) and chronological age (C and F), for each cohort UCL (Top: A-C) and Aberdeen (Bottom: D-F). P-values: (*)<0.1, *<0.05, **<0.01, ***<0.001, ****<0.0001 in the violin plots and numerically in the scatterplots.

Among the SCZ cases clozapine treatment was associated with a reduced OMR burden in the Aberdeen cohort (Figure 1B and E; Supplementary Tables GSE84727 Tab 2) and in the metaanalysis (Table 2). Chronological age was associated with an increased OMR burden in both cohorts independently (Figure 1C and F) and in the meta-analysis (Table 2). These results remained significant when retaining the outlier samples (Supplementary Figure S10 and S11 and Supplementary Tables GSE80417 and GSE84727 Tab 2).

Stratifying OMRs by direction did not reveal enrichment of either hyper- or hypo-methylated OMRs. SCZ, clozapine treatment and chronological age were significantly associated with both positive and negative OMRs (Supplementary Tables GSE80417, GSE84727, Meta Tab 3).

### Moderately deviating OMRs are enriched in SCZ cases

Stratifying the OMRs by their overall deviation from the norm (i.e. |MeanZ|) revealed that the increased OMR burden in SCZ cases was driven by moderately deviating OMRs. SCZ and clozapine treatment were associated with enrichment/depletion of OMRs deviating by 3-7 SDs, and chronological age with OMRs deviating by 3-8 SDs (see Supplementary Tables GSE80417, GSE84727 and Meta Tab 4).

These moderately deviating OMRs encompass the majority of OMRs (>90%) and more than one third of the samples have no OMRs in more extreme deviation bins. To mitigate this zero-inflation, we performed burden testing for the most deviating OMRs in a single bin (>8 SDs), b**ut** found no significant associations with SCZ or any of the covariates - despite this bin containing more OMRs than the 6-7 and 7-8 SD bins alone (see Supplementary Tables GSE80417, GSE84727 and Meta Tab 4). We also stratified the OMRs by their size (genomic coverage in bp), but found that SCZ, clozapine treatment and chronological age to be associated with the OMR burden across all size bins in both cohorts (see Supplementary Tables GSE80417, GSE84727 and Meta Tab 5).

### OMRs in SCZ patients are enriched in regulatory features

We found that OMRs in SCZ cases were not enriched in any genomic feature after the FDR correction and covariate adjustment in the individual cohorts nor in the meta-analysis (Figure 2A; Supplementary Tables GSE80417, GSE84727, meta Tab 6; Supplementary Figures S12-14).

**Figure 2:**
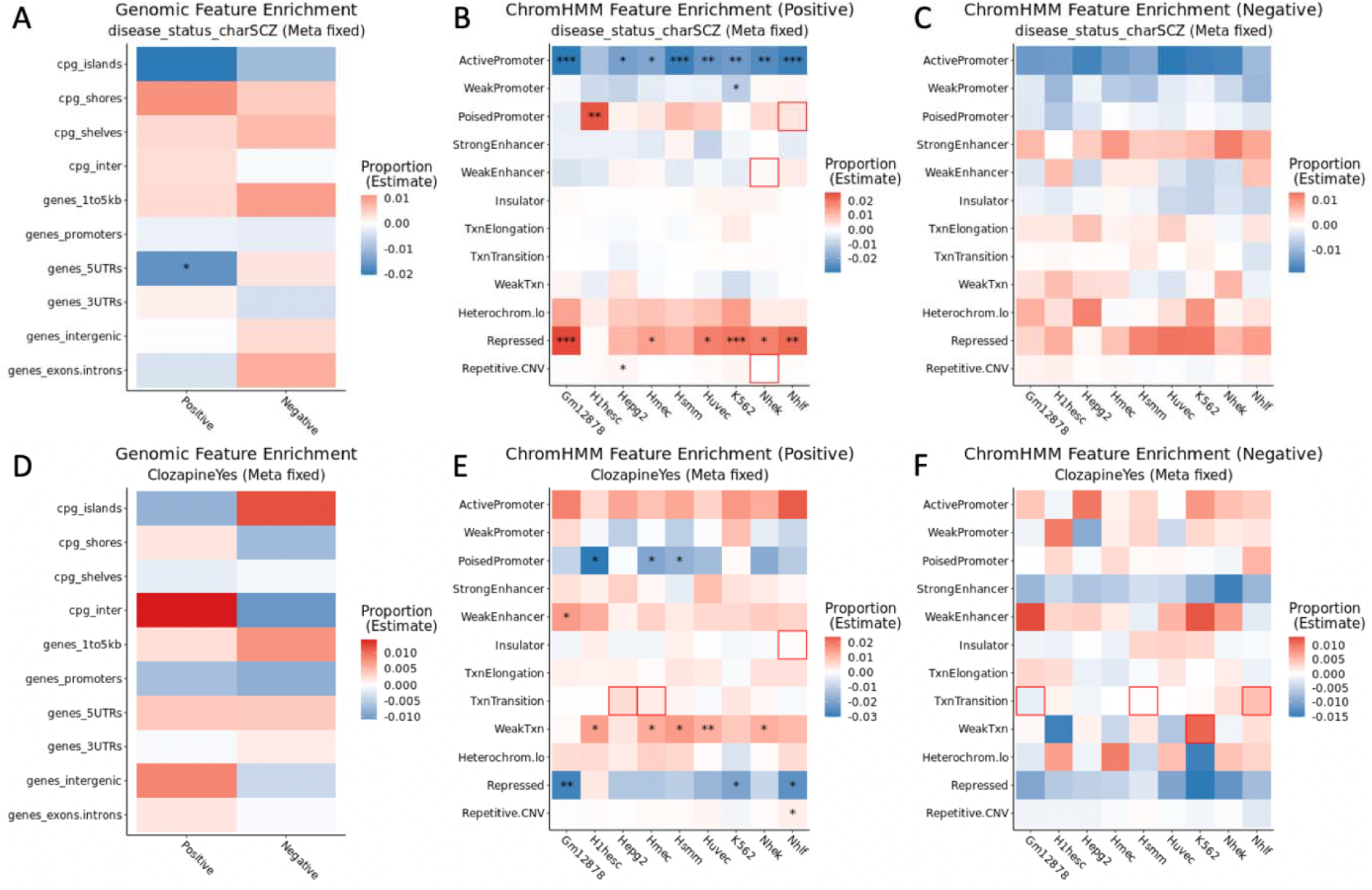
Heatmaps of enrichment of OMRs across genomic (A and D) and ChromHMM (B-C, E-F) features associated with SCZ status (top panel) and clozapine use (bottom panel). Each cell in the heatmap illustrates association with annotations derived from one of the 9 cell lines (blue = depletion, red = enrichment). All estimates and P-values are obtained from the meta-analysis of the results from the adjusted models: ~ SCZ + DNAmSS + Age (A-C), ~ SCZ + DNAmSS + Age + clozapine (D-F). Estimates and p-values are obtained from the fixed effect metaanalysis of the two cohorts. P-values: *<0.05, **<0.01, ***<0.001, ****<0.0001, and hetP<0.05 is indicated in red boxes.

However, the ChromHMM epigenomic tracks revealed hyper-methylated (positive) OMRs in SCZ cases were enriched in polycomb *repressed* elements in several non-stem cell lines, and in *poised promoters* in H1hesc stem cell line (Figure 2B; Supplementary Figures S18-20 and Supplementary Tables GSE80417, GSE84727, Meta Tab 7). This was accompanied by a relative depletion in *active promoters* across most cell lines. These effects remained significant after adjustment for covariates, and we found ageing to independently associate with similar regions (Supplementary Figure S21). Enrichment patters associated with clozapine treatment were often opposite of those associated with SCZ (Figure 2E; Supplementary Figure S21-23 and Supplementary Tables GSE80417, GSE84727 and Meta Tab 7). Clozapine treatment was also independently associated with an enrichment of OMRs in weak *enhancers* and *transcribed* regions.

Hypo-methylated (negative) OMRs showed no enrichment in association with SCZ status or clozapine treatment (Figure 2C and F). Associations with covariates are shown in Supplementary Figures S27-29.

### Individuals with OMRs in developmental gene sets are enriched for SCZ cases

Genic OMRs included over 80% of all OMRs in each cohort. Individuals with enrichment of OMRs in *developmental processes* were more likely to be SCZ cases (meta-analysis OR [95%CI] = 2.37 [1.56-3.62], FDR=0.003), and this remained significant after controlling for the covariates (see Supplementary Tables Meta Tab 9).

Since detection of the overrepresented pathways may be confounded by the OMR burden, we also tested if SCZ cases were enriched for having *“any”* overrepresented pathway (i.e. nonspecific overrepresentation) as well as pathways not assigned to any category (“misc” category). We found individuals having *any* pathway to be enriched for SCZ cases, but was less significant than the specific association, and we cannot rule out that these findings may be confounded by the increased OMR burden (see Supplementary Tables GSE80417, GSE84727 and Meta Tab 9).

## Discussion

We used an optimization algorithm to detect and quantify individual-level epigenetic deviation in genome-wide DNAm array data – which we call outlier methylation regions (OMRs). The optimization ensures detection of similarly deviating, consecutive probes - making them unlikely to arise due to technical artifacts, and facilitates their quantitative examination (i.e. direction, extent of deviation from the norm, size and genomic context).

We used two independent, publicly available case-control SCZ cohorts, and in both datasets we found a highly significant and comparable increase in the overall burden of OMRs in SCZ cases. This suggests a potential role of epigenetic dysregulation – in the form of private/rare events – in SCZ pathology, in addition to the case-control group differences that have previously been assessed in these datasets ^14^. Importantly, we also observed lower burden of OMRs in clozapine-treated, compared to untreated SCZ cases. In the paragraphs below we discuss the potential relevance of these findings to SCZ etiology, and the caveats warranting caution in their interpretation.

In order to characterize the OMRs enriched in SCZ cases, we assessed if they differed with respect to size, direction and deviation compared to the OMRs identified in the controls. We found that the SCZ cases had a higher burden of moderately deviating OMRs, but found no evidence for an increased burden of the most deviating (> 8SDs), and very rare OMRs – despite the total OMR count in this extreme bin being higher than the OMR count in the 6-7 and 7-8 SDs bins, indicating that the lack of significance was not due to lack of power to detect casecontrol differences in sparse OMR bins. This is in line with a previous study in the same datasets, using a different method to detect extremely deviant and rare OMRs (i.e. *epivariations*), which found no overall burden increase in SCZ ^37^. The additional functional insights derived from the analyses of the moderately-deviating OMRs in our study highlight the utility of systematic detection followed by stratification of OMRs at different deviations (from low to high). We observed no evidence that the OMR enrichment in SCZ cases is underlain by events of certain size or direction.

Annotating OMRs according to their (epi)genomic context, we observed remarkable similarities between the two cohorts. The OMRs in SCZ cases were significantly enriched in poised promoters in stem cells, and in polycomb repressed elements in differentiated cell lines - features that indicate impacts on cellular differentiation ^38^ and somatic development ^39^. This is in line with a previous (EWAS) finding of DNAm differences in adult prefrontal cortex in regions associated with repressive chromatin marks ^40^. Our results, showing an increased number of SCZ cases having genic OMRs overrepresented in developmental pathways, further support that OMRs may impact, or be caused by, alterations in the developmental processes. Collectively, these results point towards epigenetic dysregulation of developmental processes in SCZ, and are consistent with a developmental origin of SCZ.

Clozapine treatment exhibited a reverse pattern of OMR enrichment in SCZ cases (i.e. depletion specifically in poised promoters and polycomb repressed regions; NB. unlike in the main analyses, the controls were non-treated SCZ cases, rather than unaffected individuals), but was also associated with an enrichment of OMRs across weak enhancers and transcribed regions. The latter finding is in line with clozapine’s profound impact on the methylome ^41^, but whether clozapine also “normalizes” the methylome in treated SCZ cases is not clear. Clozapine is an atypical antipsychotic given to patients with severe, treatment-resistant SCZ - a potentially unique phenotype, and the reduced OMR burden found in our study may stem from underlying phenotypic differences, or alternative medication use.

Aging was also associated with increased OMR burden and enriched in ChromHMM features similar to SCZ for positive OMRs and more for negative OMRs. While only the UCL cohort was skewed towards older cases than controls, we cannot fully rule out a confounding impact, despite adjustment in our models.

The exact role of the OMRs in SCZ remains to be established. An increased burden of OMRs in SCZ cases can reflect downstream effects of rare genetic risk variants, environmental insults potentially leading to, or correlated with disease, or simply be caused by the disease/medication itself (i.e. reverse causation). In favor of the former, a recent study demonstrated an increased burden of structural variants affecting the chromatin 3D structure in SCZ ^7^, which could possibly underlie large-scale methylomic dysregulation in SCZ cases. Nevertheless, causal processes underlying disease cannot be unequivocally inferred from adult epigenetic data – as changes in DNAm observed in cases cannot distinguish causes from consequences of having the disease, e.g. receiving medications ^42^. Secondly, we could not assign an independent genetic or environmental origin to the OMRs in SCZ, nor establish whether their increased burden in blood of SCZ cases would replicate in other tissues. To address these issues, our findings warrant further studies in longitudinal and twin study settings, and/or in cohorts with complete treatment and genetic information, as well as across other tissues.

In summary, we detected and quantified rare methylomic outliers in SCZ (OMRs), and found an increased burden of functionally-distinct, methylomic events in SCZ cases compared to controls – along with a reverse pattern associated with clozapine treatment in cases. Previous studies of outlier methylation have focused on the most deviant regions (i.e. *epivariations)^16,17,43^*. We have now extended the spectrum of OMR detection by designing a new method (OMA) - to systematically detect and quantify regions with moderate to extreme deviation in DNAm levels. The application of our method is limited only by the computational demands of the optimization step - restrained by setting a lower OMR detection boundary (here 3 SD), and can be applied across different array designs. We offer OMA to the scientific community to detect and quantify OMR in their own methylomic datasets and, in addition to EWAS analysis, assess rare epigenetic events in diseases with a likely heterogeneous etiology.

## Data and code availability

DNA methylation data is available through Gene Expression Omnibus under accession numbers [GEO:GSE80417] and [GEO:GSE84727].

Code for Outlier Methylation Analysis is available at: https://gitlab.com/CSoeholm/outlier-methylation-analysis-oma

