## Supplemental Figures for "Assessing the burden of rare DNA methylation deviations in schizophrenia": SupplementaryFigures.pptx

### Slide 1
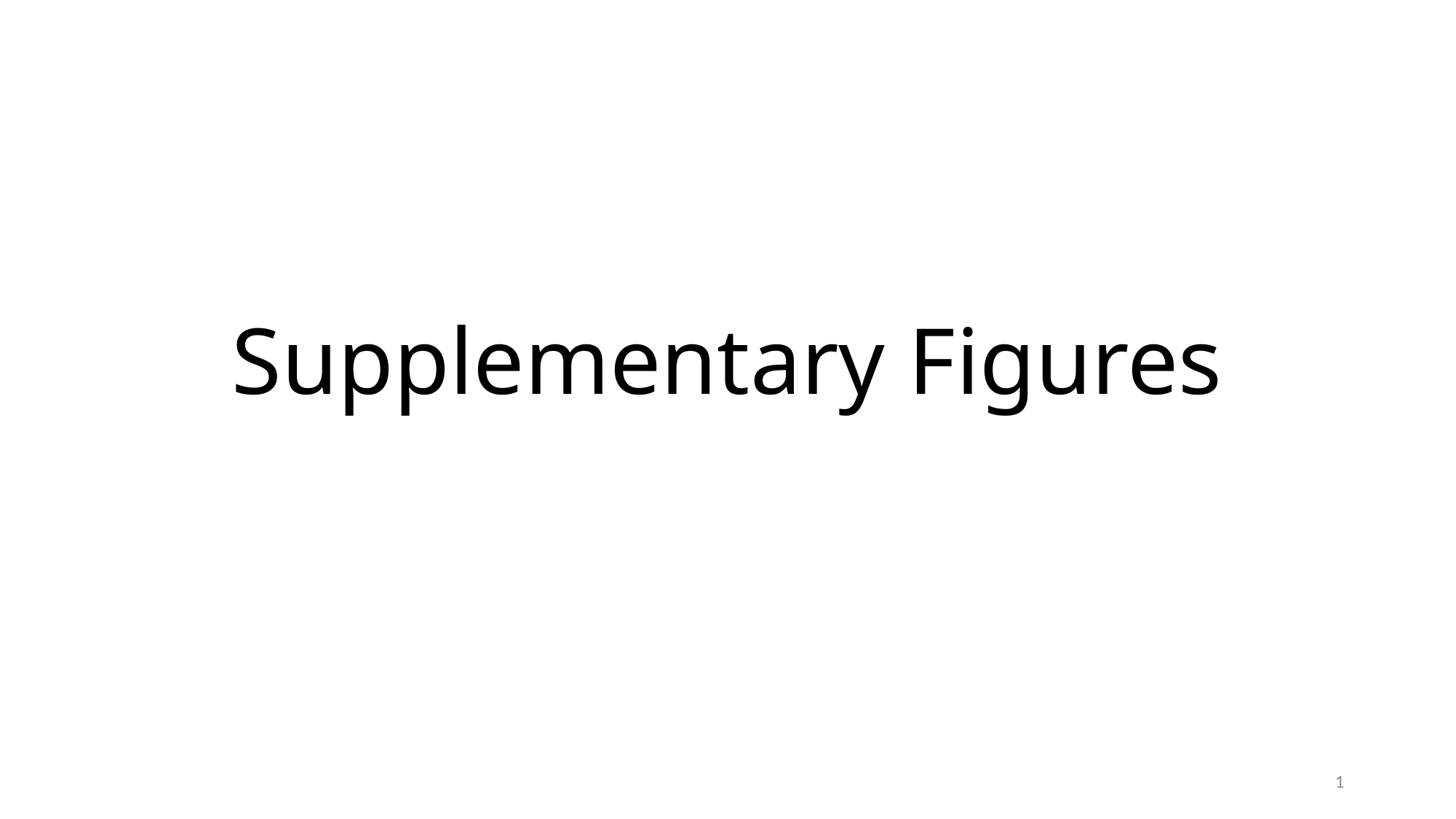

#### Supplementary Figures
1

### Slide 2
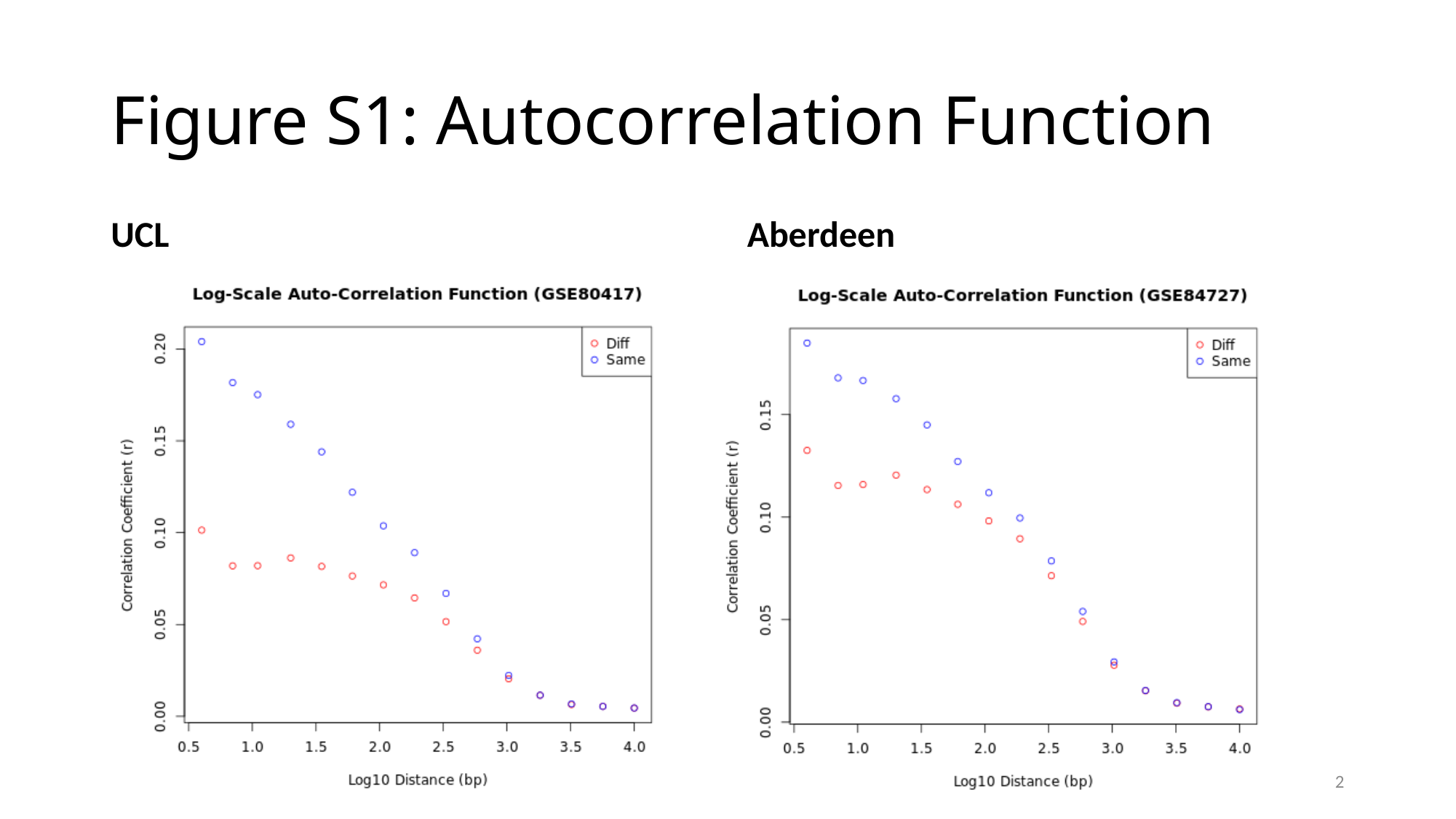

#### Figure S1: Autocorrelation Function
UCL
Aberdeen
2

### Slide 3
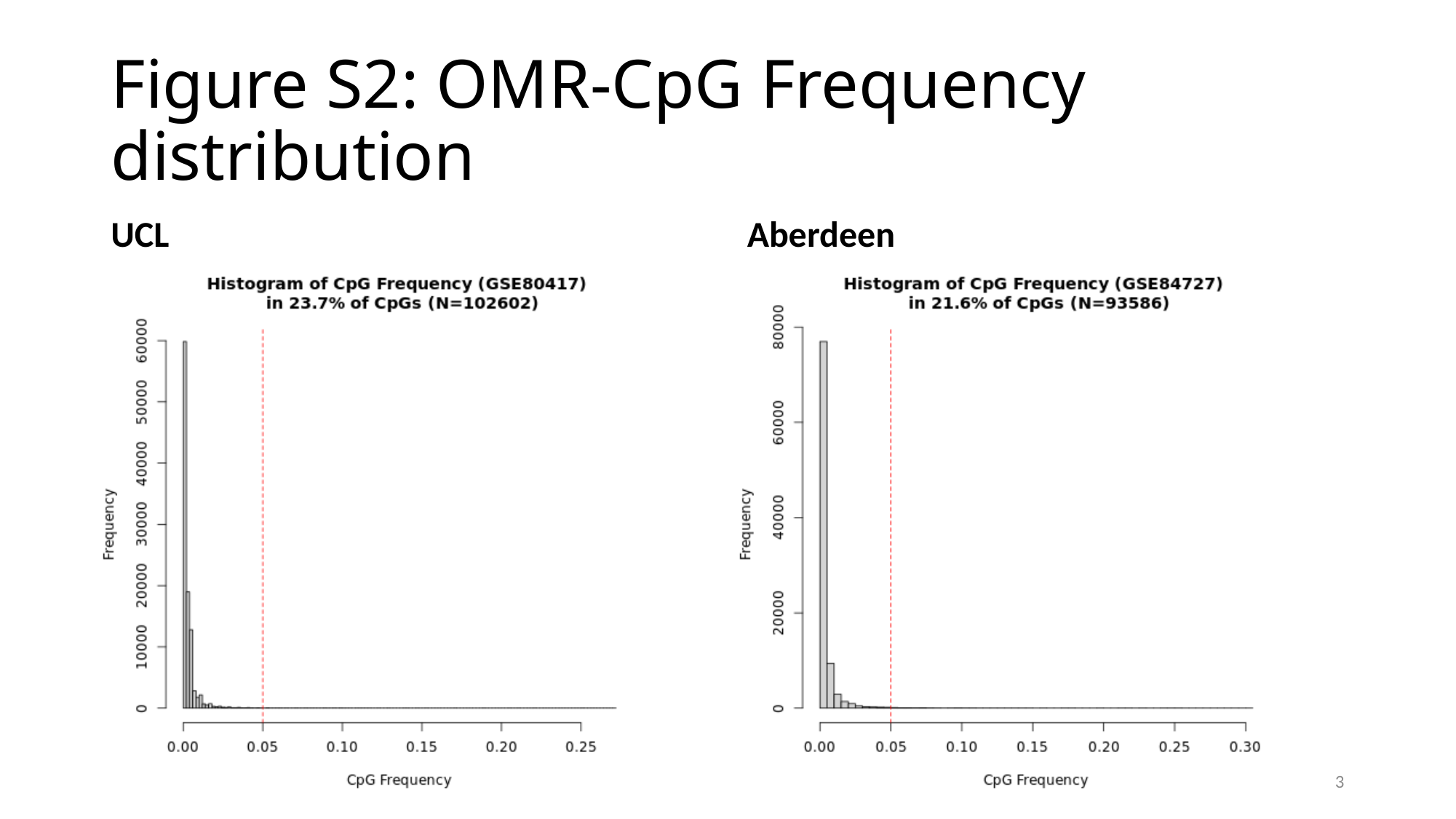

#### Figure S2: OMR-CpG Frequency distribution
UCL
Aberdeen
3

### Slide 4
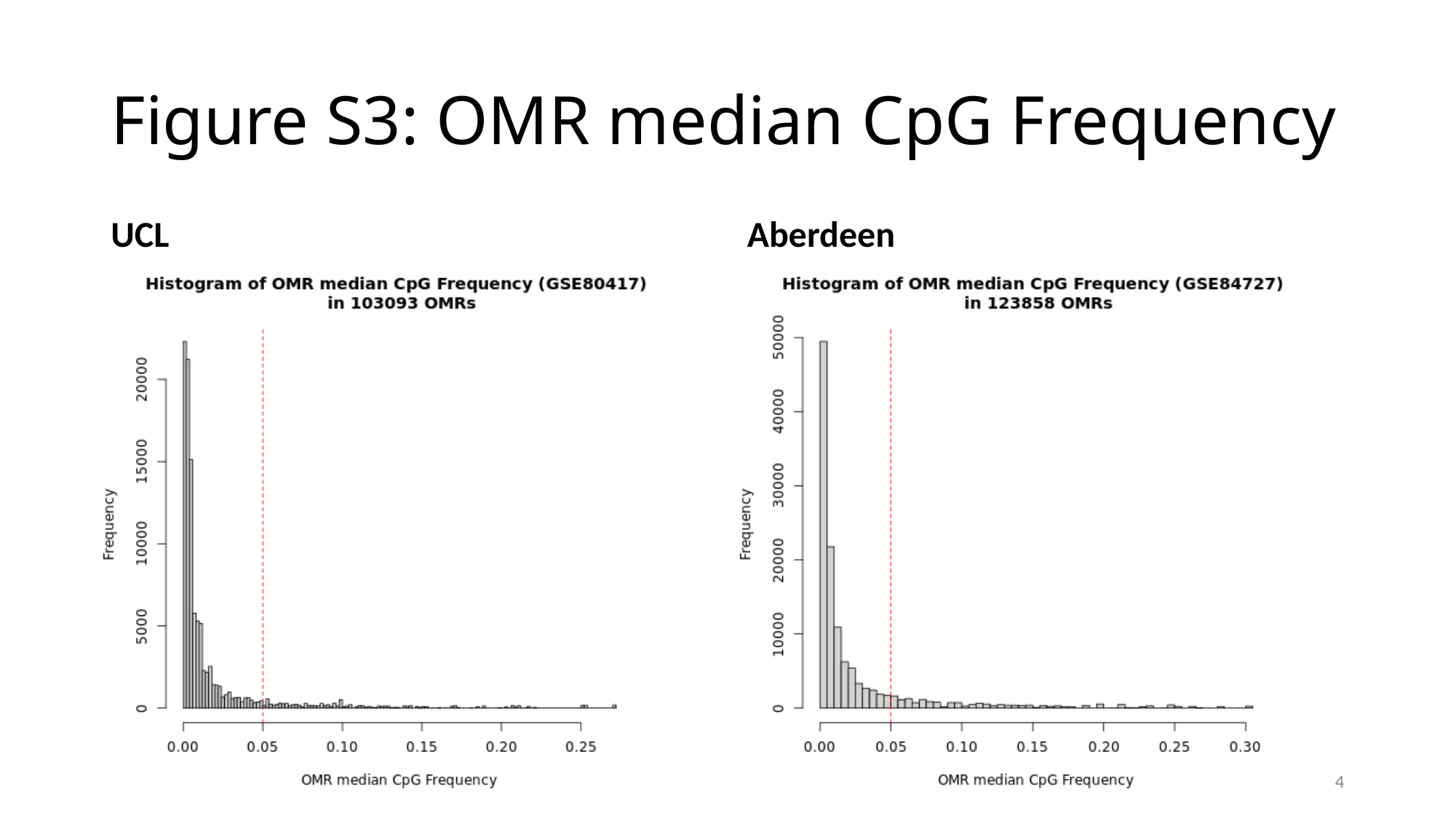

#### Figure S3: OMR median CpG Frequency
UCL
Aberdeen
4

### Slide 5
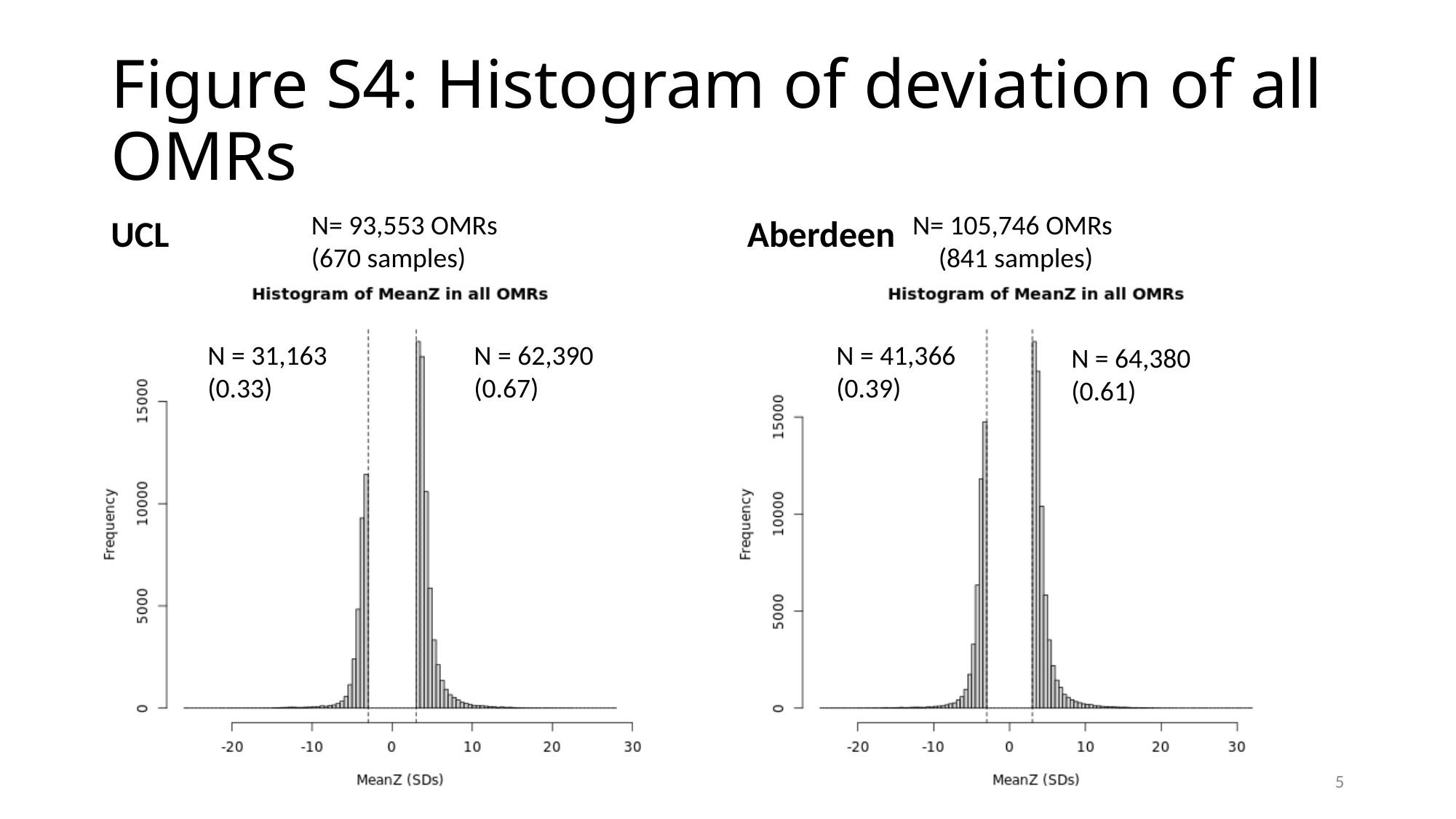

#### Figure S4: Histogram of deviation of all OMRs
UCL
Aberdeen
N= 93,553 OMRs
(670 samples)
N= 105,746 OMRs
(841 samples)
N = 41,366
(0.39)
N = 31,163
(0.33)
N = 62,390
(0.67)
N = 64,380
(0.61)
5

### Slide 6
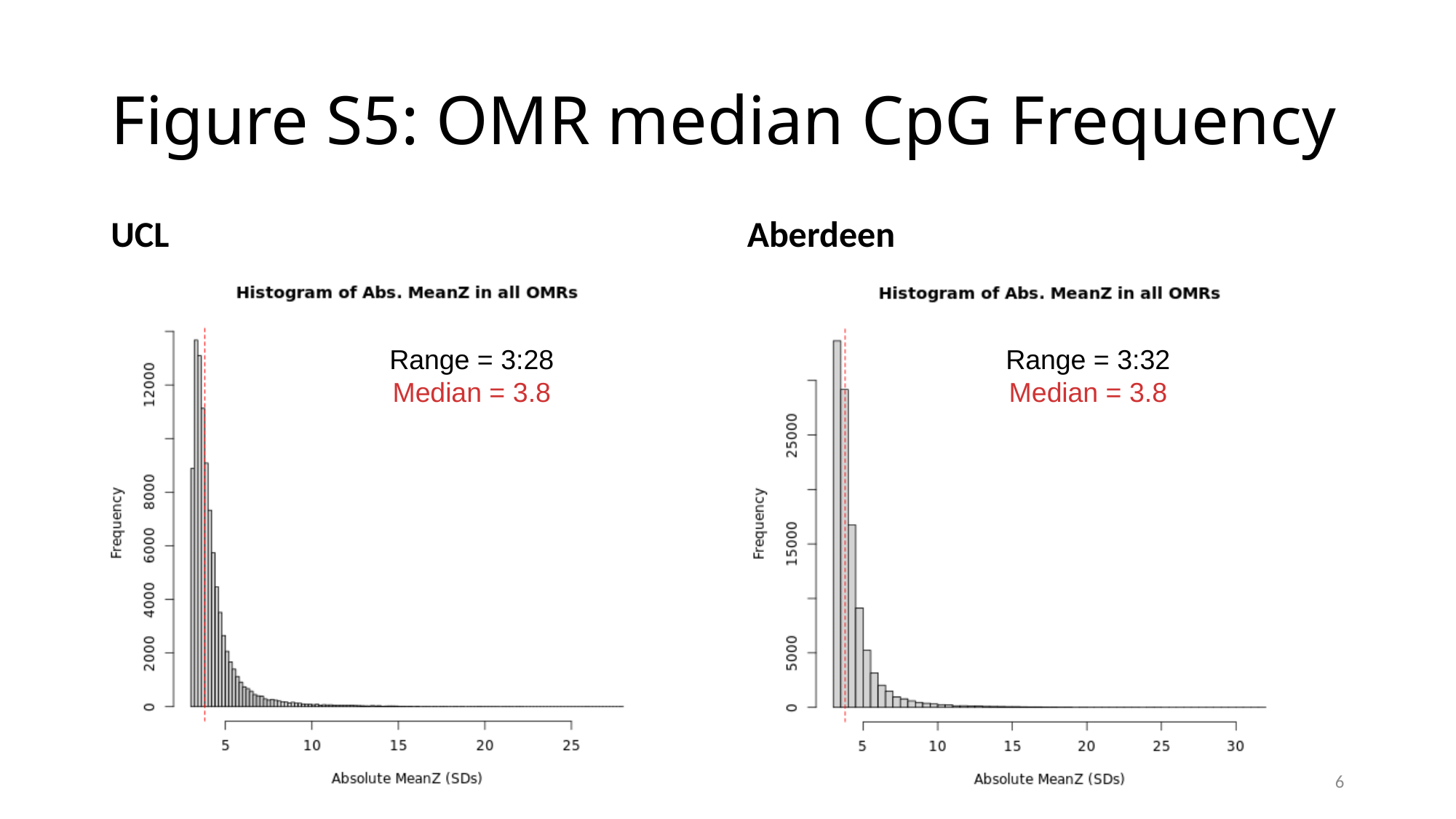

#### Figure S5: OMR median CpG Frequency
UCL
Aberdeen
Range = 3:28
Median = 3.8
Range = 3:32
Median = 3.8
6

### Slide 7
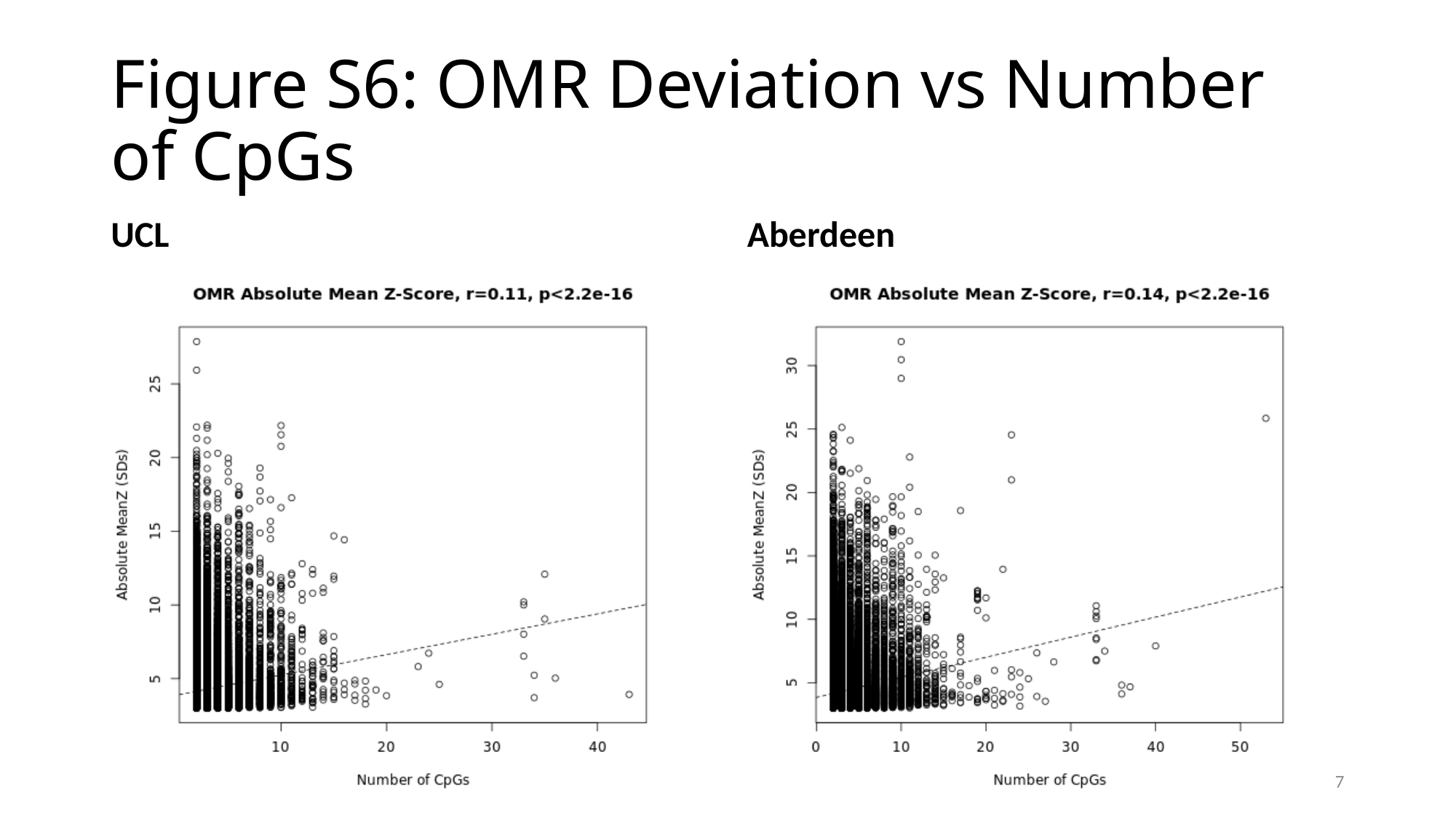

#### Figure S6: OMR Deviation vs Number of CpGs
UCL
Aberdeen
7

### Slide 8
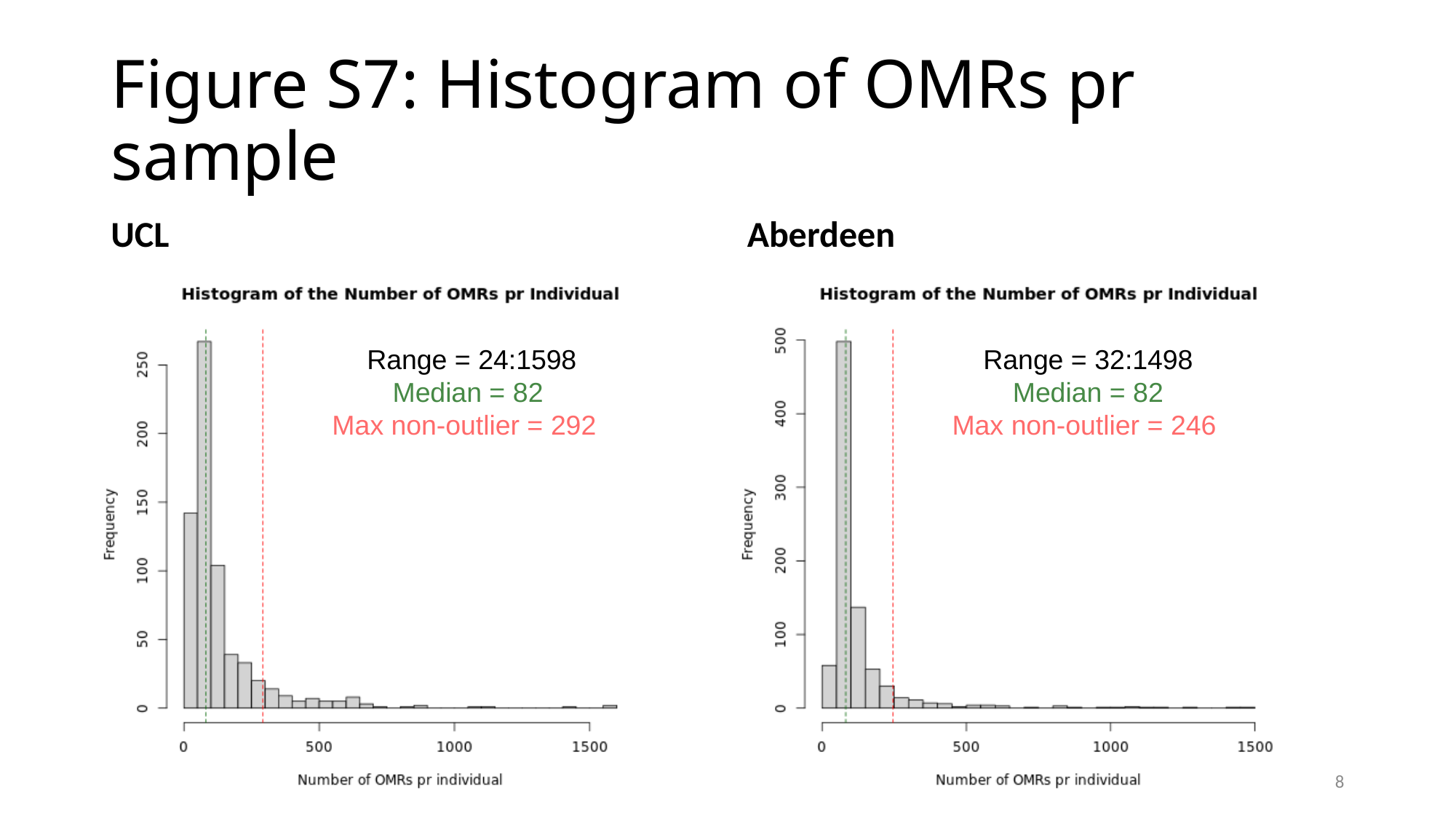

#### Figure S7: Histogram of OMRs pr sample
UCL
Aberdeen
Range = 24:1598
Median = 82
Max non-outlier = 292
Range = 32:1498
Median = 82
Max non-outlier = 246
8

### Slide 9
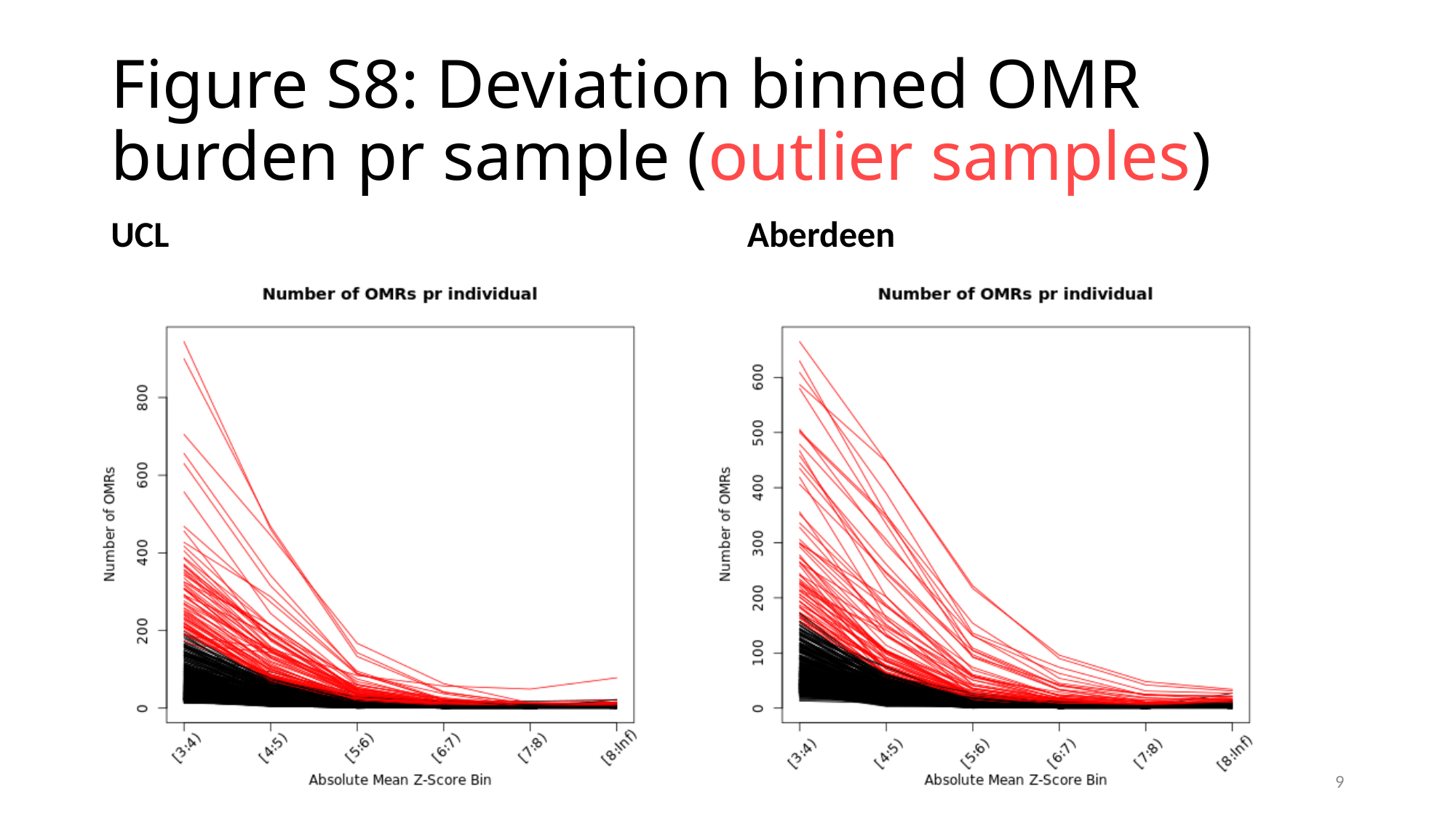

#### Figure S8: Deviation binned OMR burden pr sample (outlier samples)
UCL
Aberdeen
9

### Slide 10
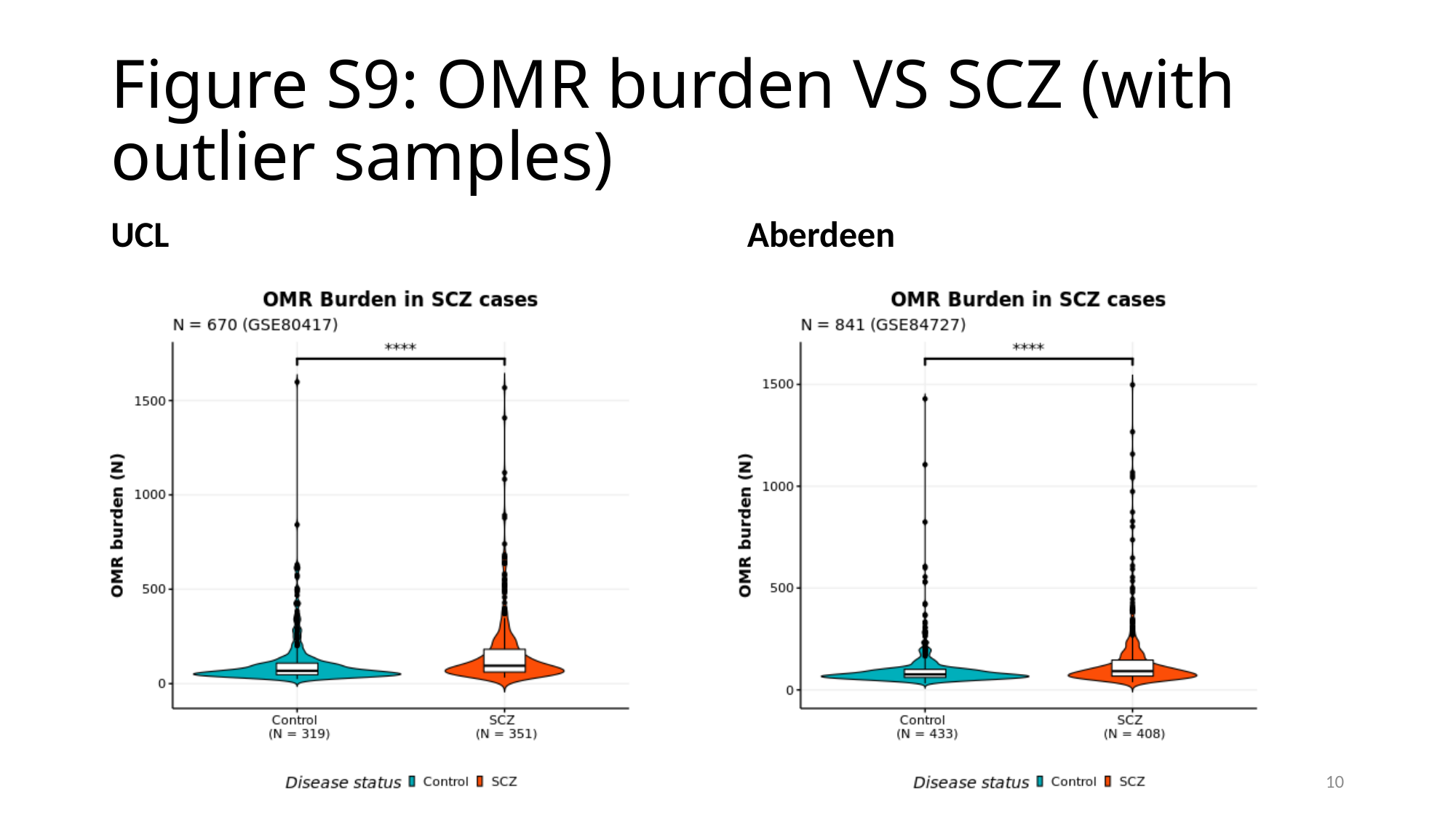

#### Figure S9: OMR burden VS SCZ (with outlier samples)
UCL
Aberdeen
10

### Slide 11
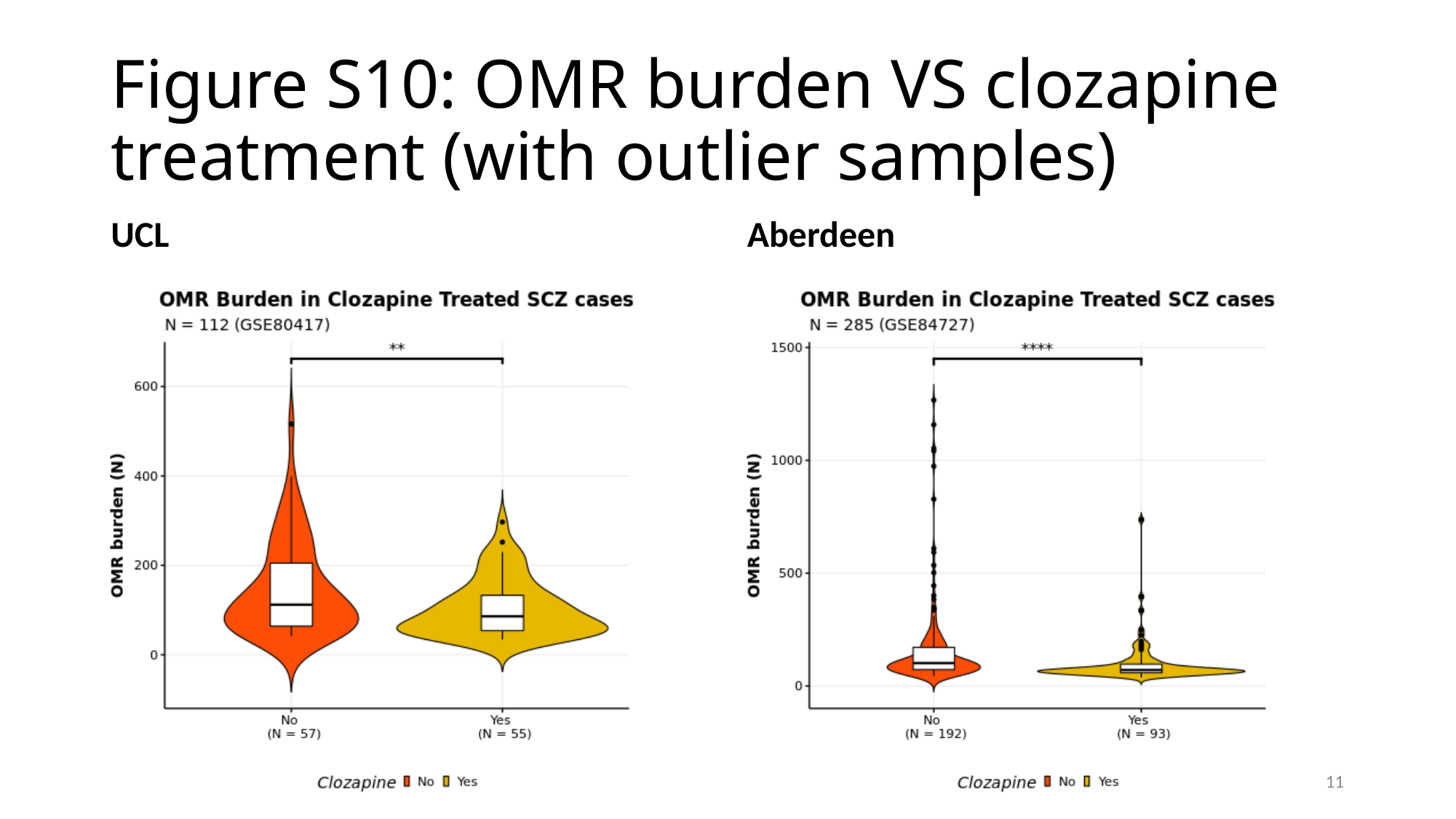

#### Figure S10: OMR burden VS clozapine treatment (with outlier samples)
UCL
Aberdeen
11

### Slide 12
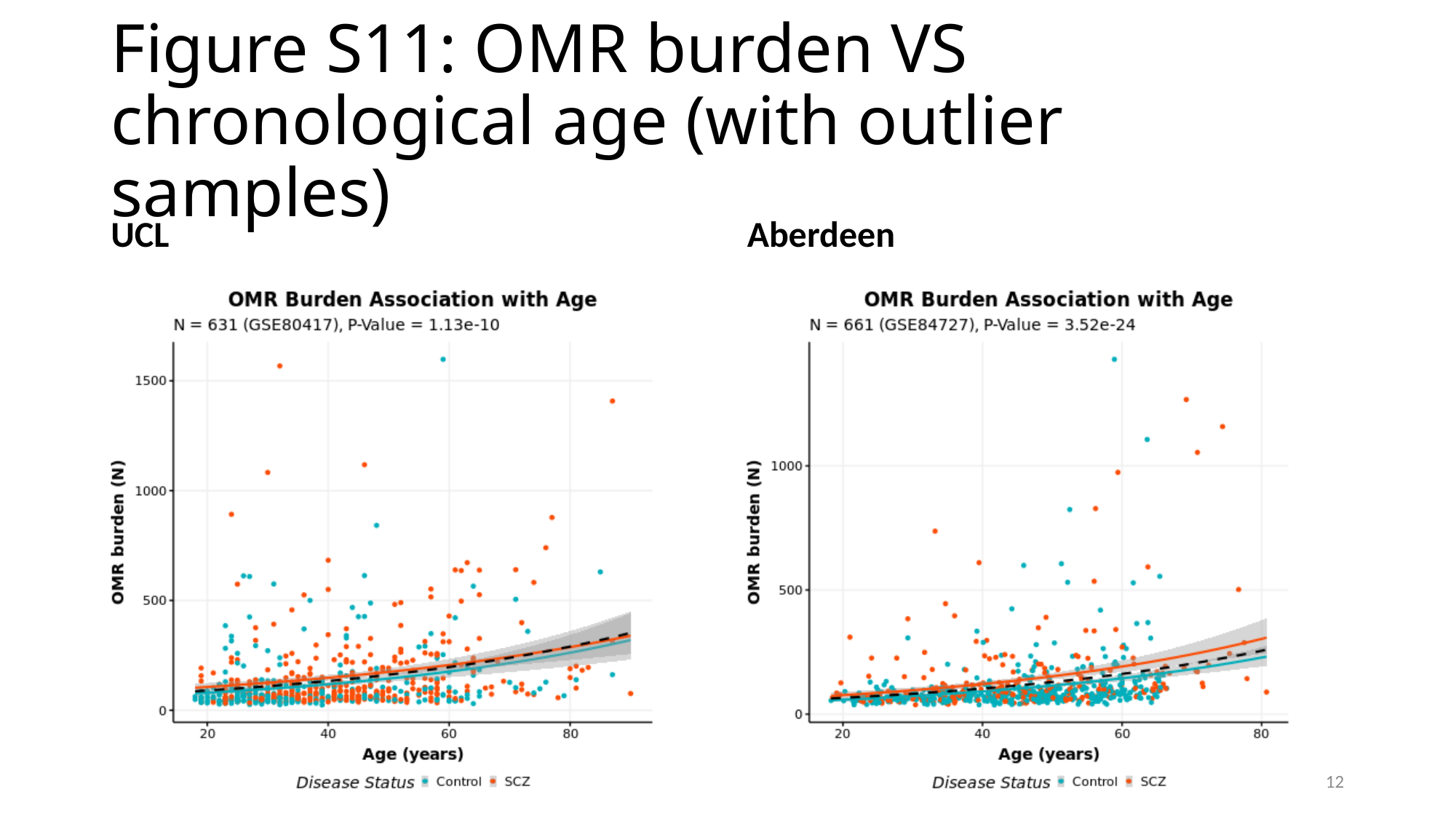

#### Figure S11: OMR burden VS chronological age (with outlier samples)
UCL
Aberdeen
12

### Slide 13
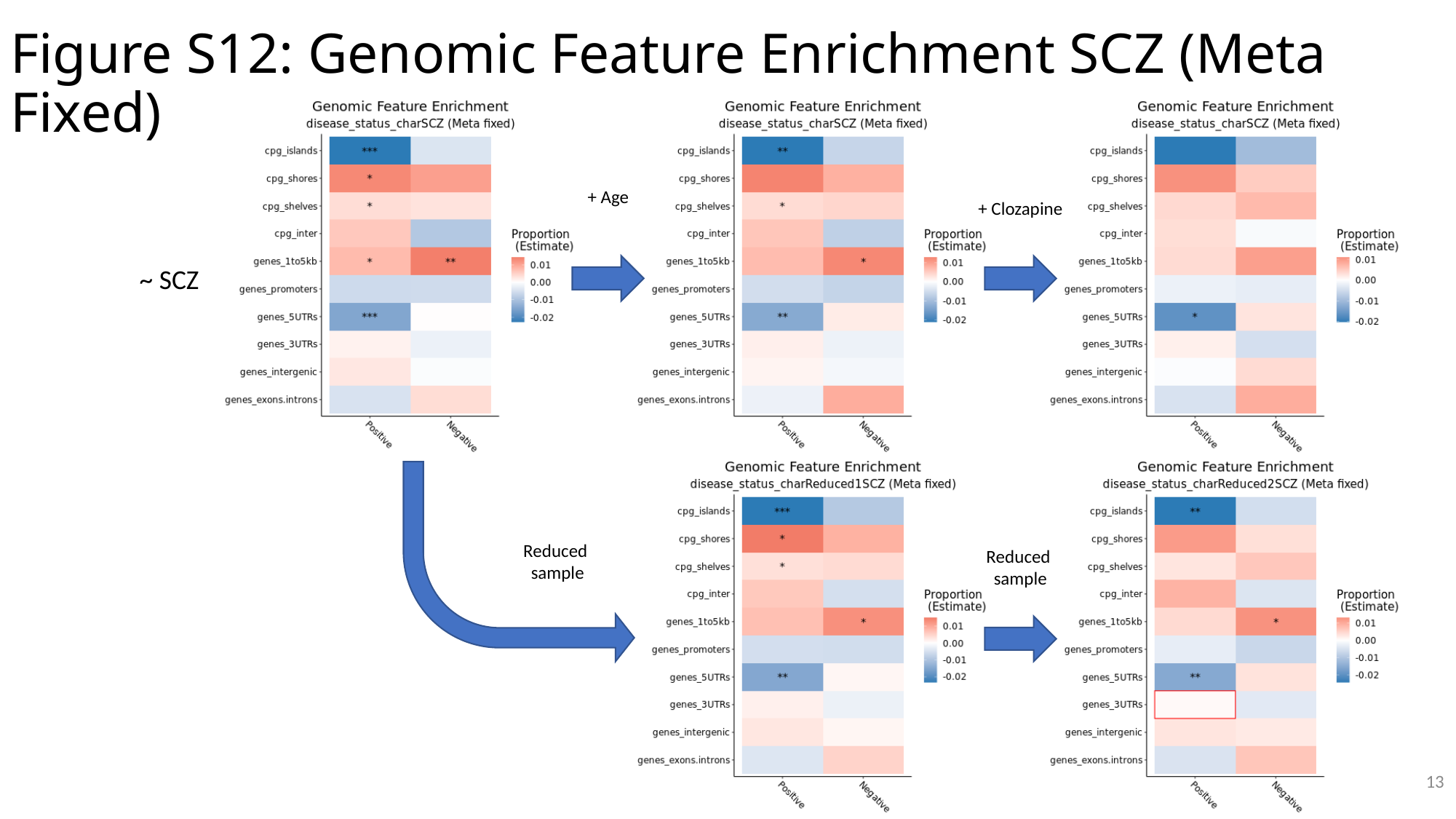

Figure S12: Genomic Feature Enrichment SCZ (Meta Fixed)
+ Age
+ Clozapine
~ SCZ
Reduced
sample
Reduced
sample
13

### Slide 14
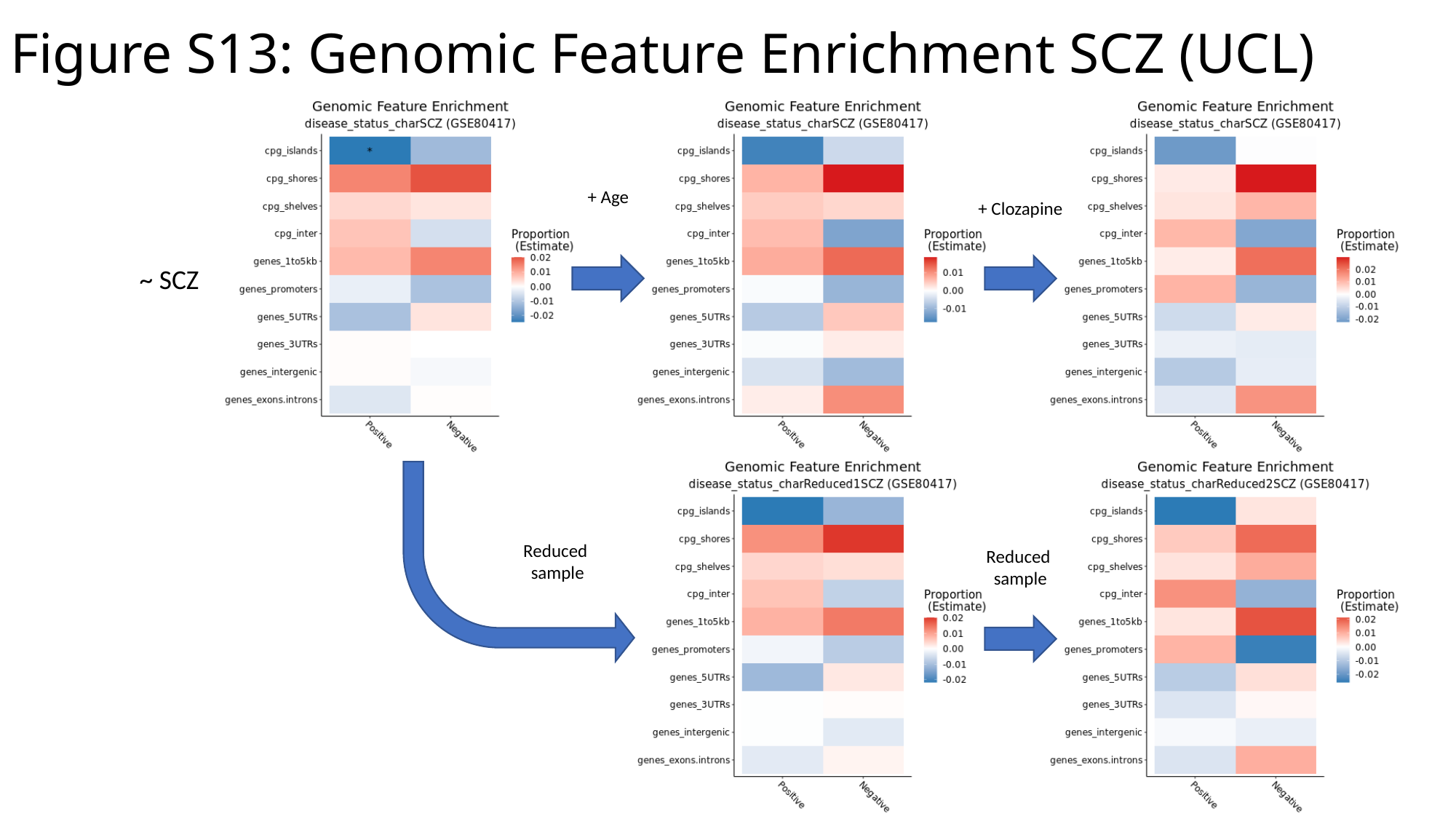

Figure S13: Genomic Feature Enrichment SCZ (UCL)
+ Age
+ Clozapine
~ SCZ
Reduced
sample
Reduced
sample

### Slide 15
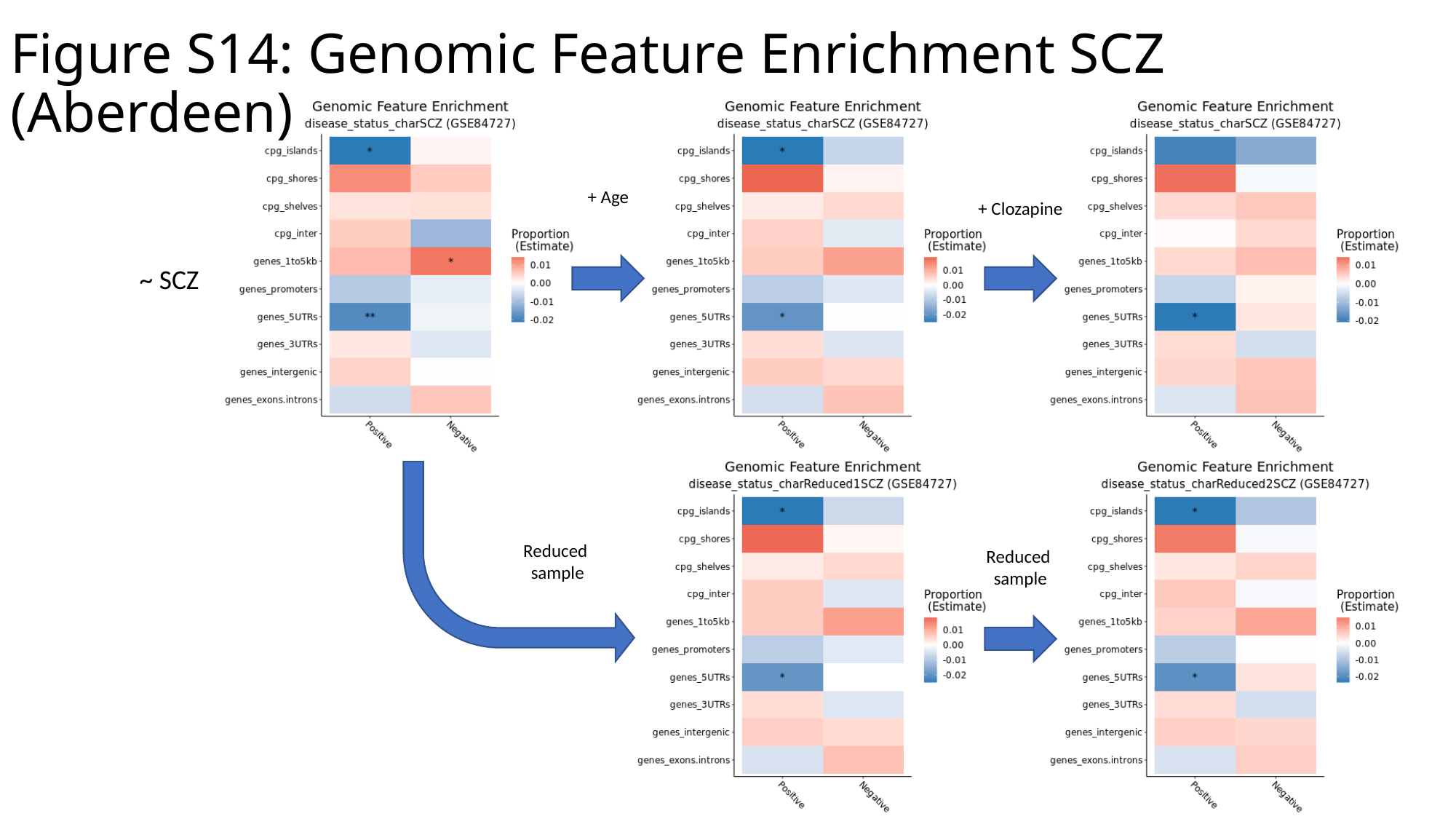

Figure S14: Genomic Feature Enrichment SCZ (Aberdeen)
+ Age
+ Clozapine
~ SCZ
Reduced
sample
Reduced
sample

### Slide 16
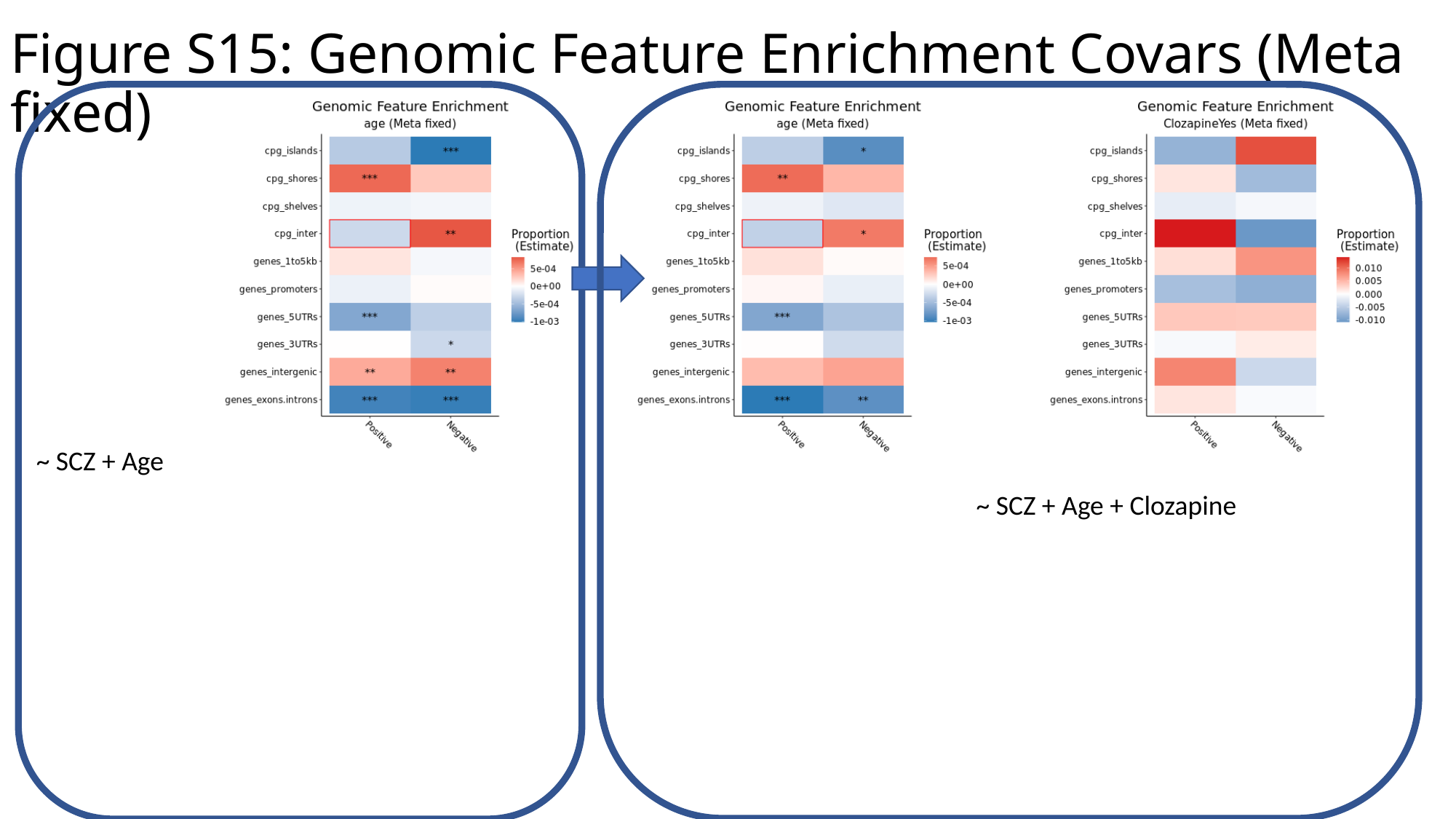

Figure S15: Genomic Feature Enrichment Covars (Meta fixed)
~ SCZ + Age
~ SCZ + Age + Clozapine

### Slide 17
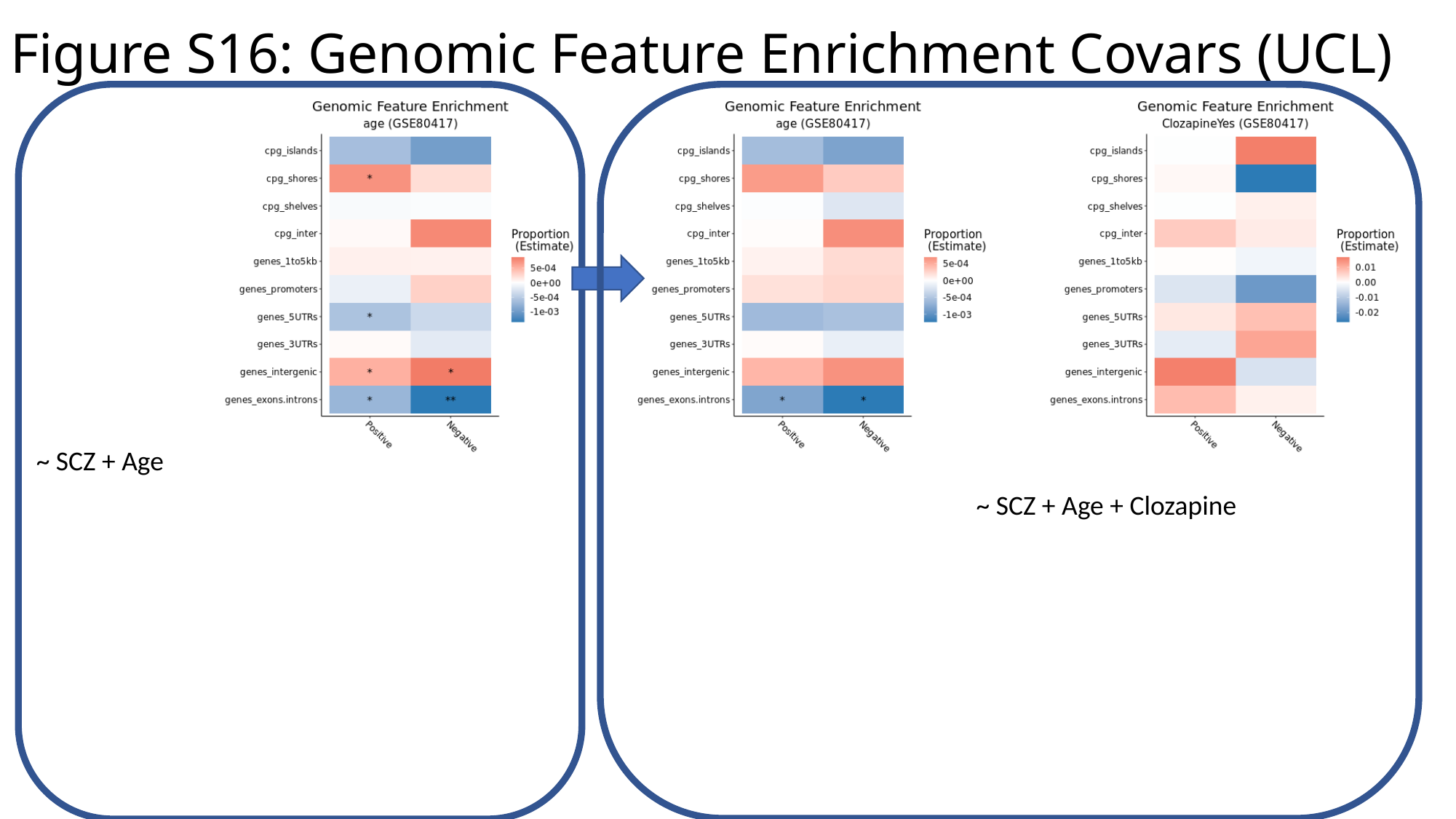

Figure S16: Genomic Feature Enrichment Covars (UCL)
~ SCZ + Age
~ SCZ + Age + Clozapine

### Slide 18
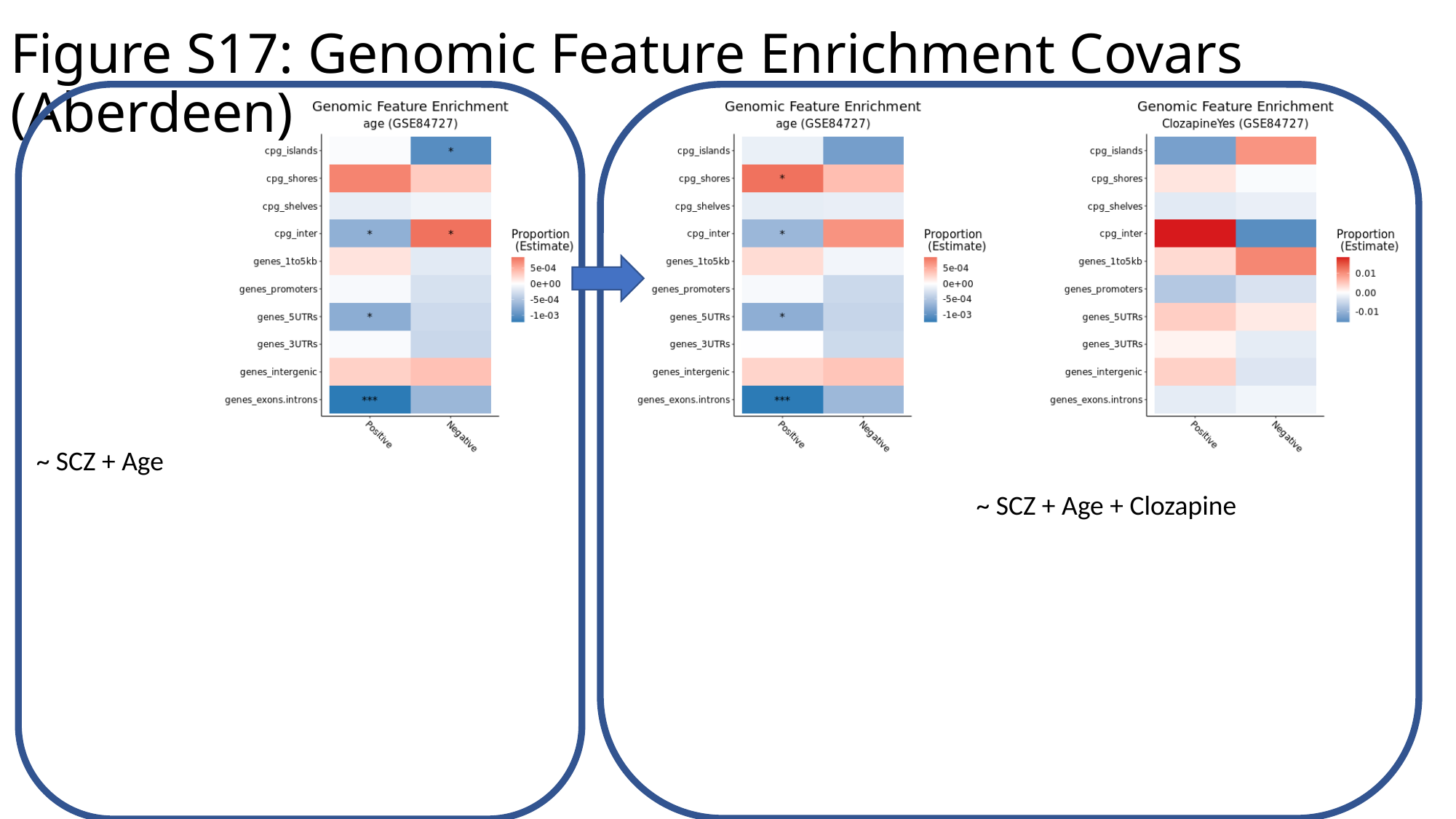

Figure S17: Genomic Feature Enrichment Covars (Aberdeen)
~ SCZ + Age
~ SCZ + Age + Clozapine

### Slide 19
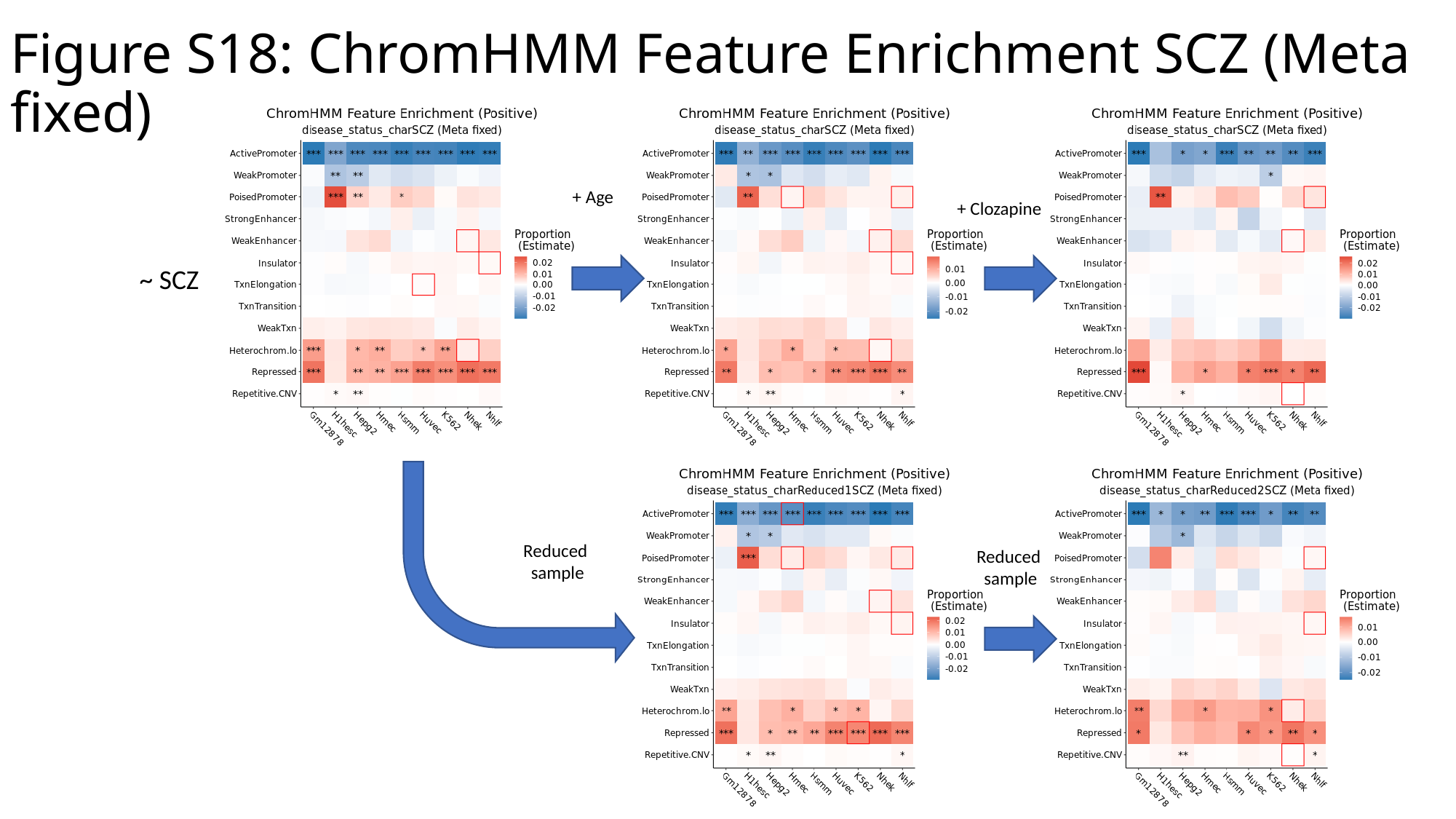

Figure S18: ChromHMM Feature Enrichment SCZ (Meta fixed)
+ Age
+ Clozapine
~ SCZ
Reduced
sample
Reduced
sample

### Slide 20
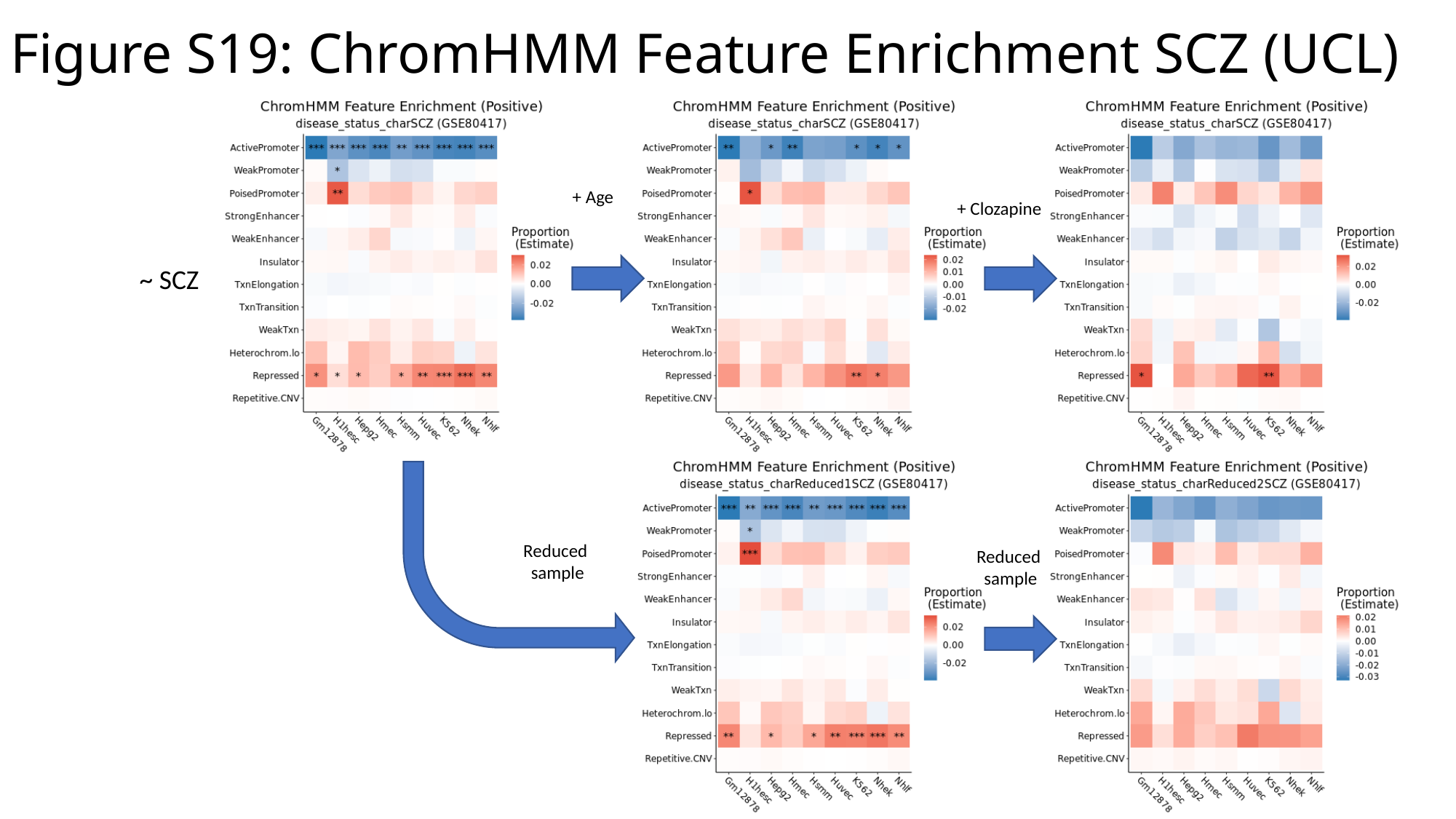

Figure S19: ChromHMM Feature Enrichment SCZ (UCL)
+ Age
+ Clozapine
~ SCZ
Reduced
sample
Reduced
sample

### Slide 21
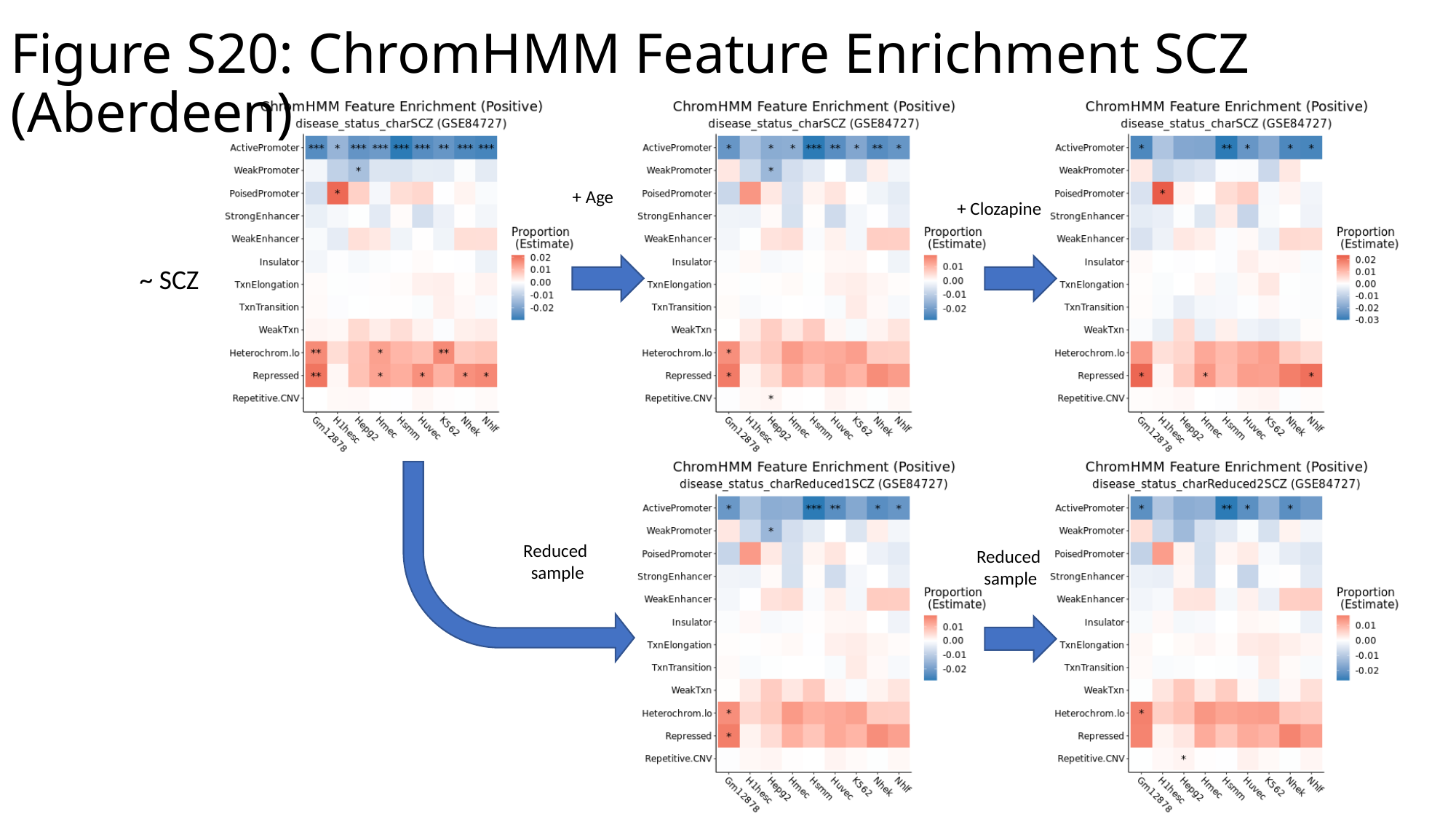

Figure S20: ChromHMM Feature Enrichment SCZ (Aberdeen)
+ Age
+ Clozapine
~ SCZ
Reduced
sample
Reduced
sample

### Slide 22
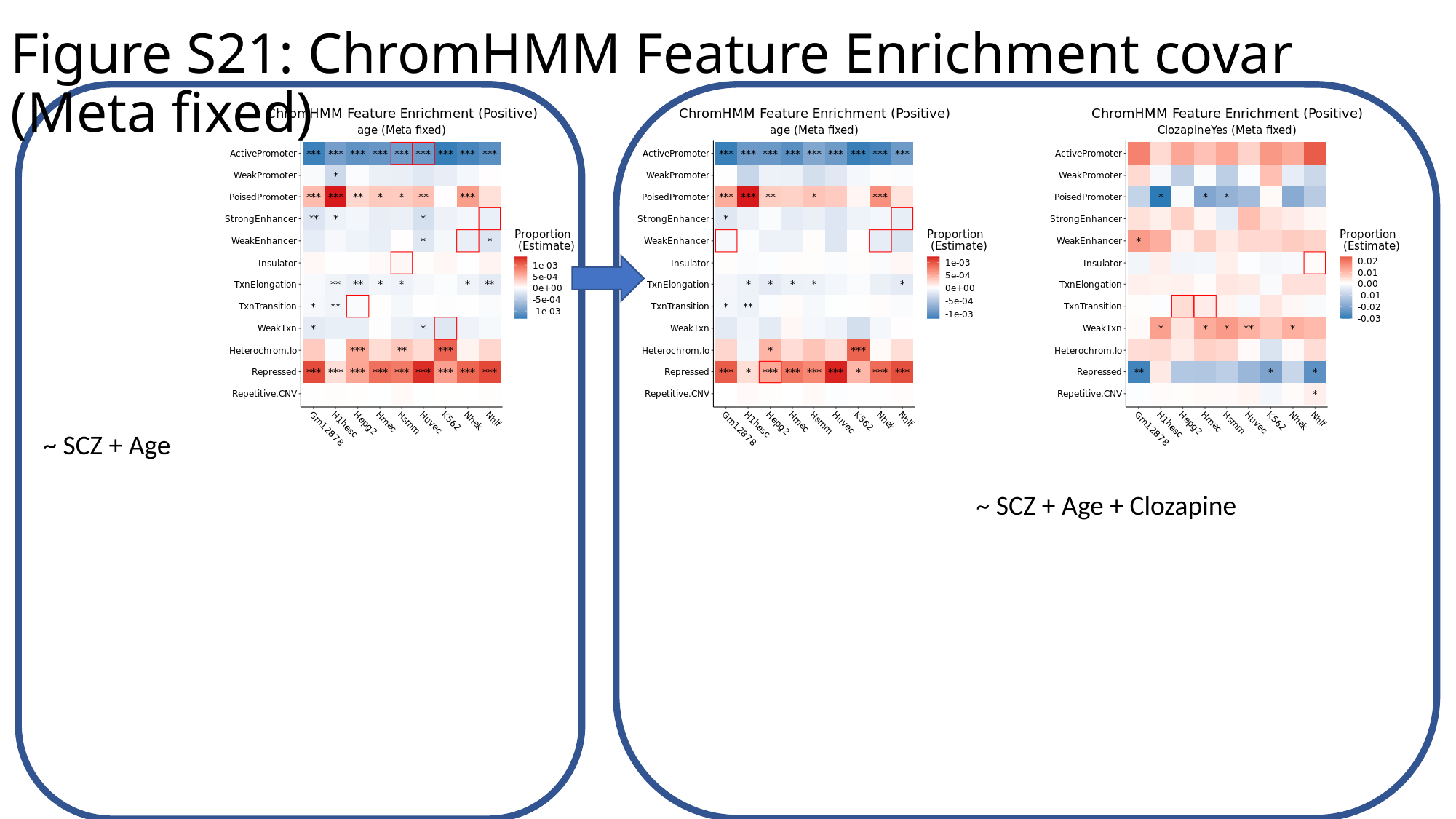

Figure S21: ChromHMM Feature Enrichment covar (Meta fixed)
~ SCZ + Age
~ SCZ + Age + Clozapine

### Slide 23
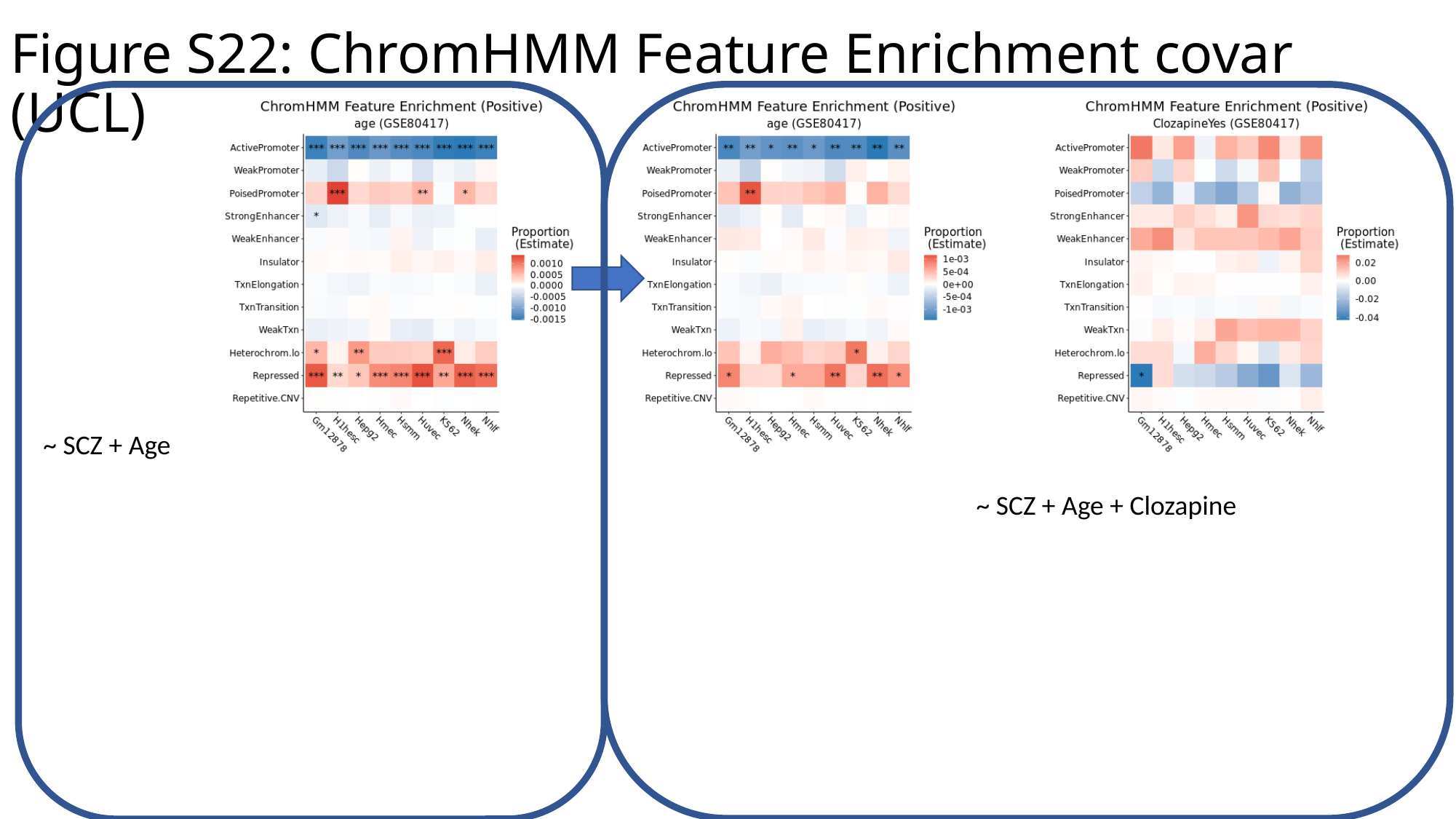

Figure S22: ChromHMM Feature Enrichment covar (UCL)
~ SCZ + Age
~ SCZ + Age + Clozapine

### Slide 24
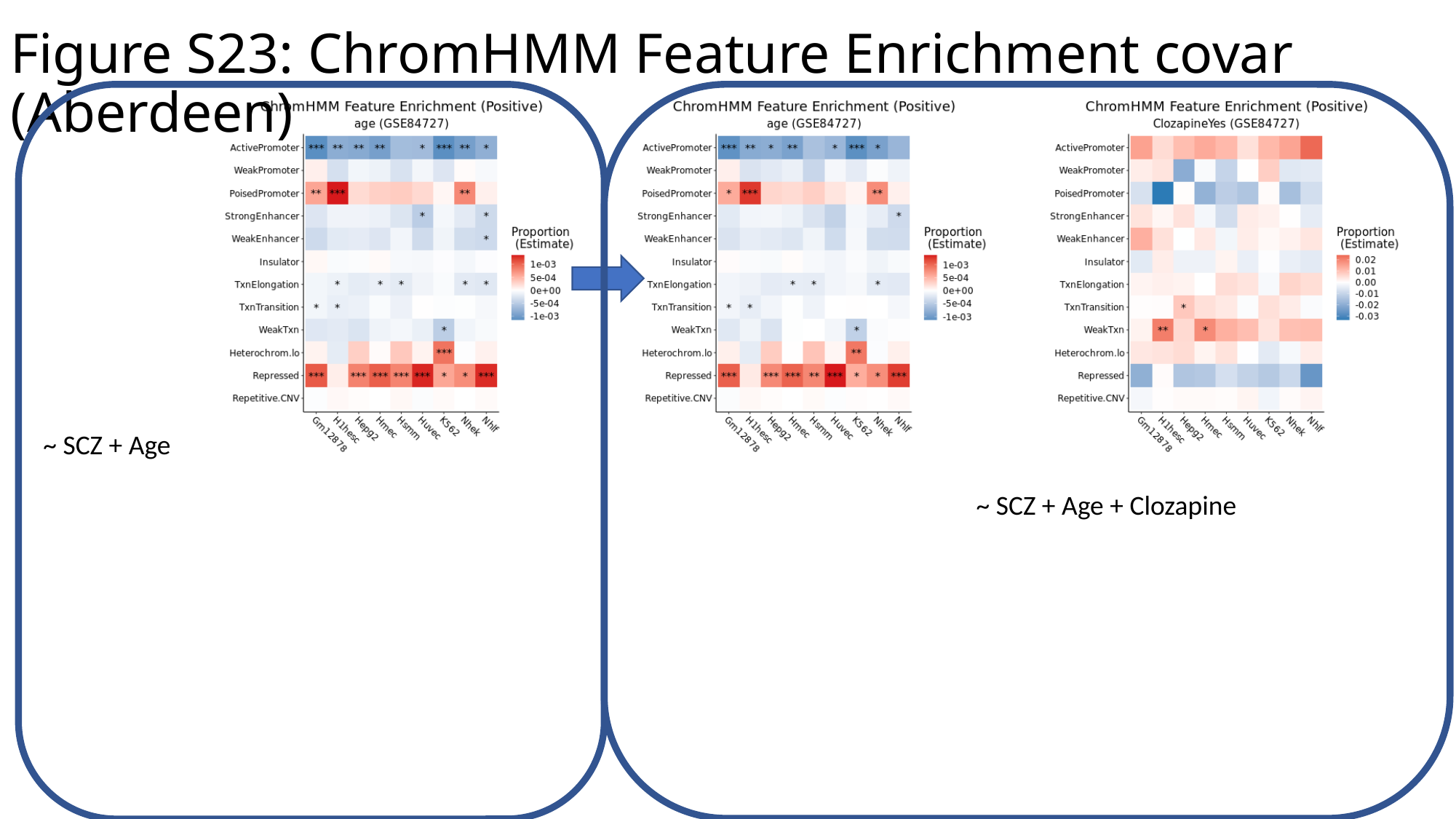

Figure S23: ChromHMM Feature Enrichment covar (Aberdeen)
~ SCZ + Age
~ SCZ + Age + Clozapine

### Slide 25
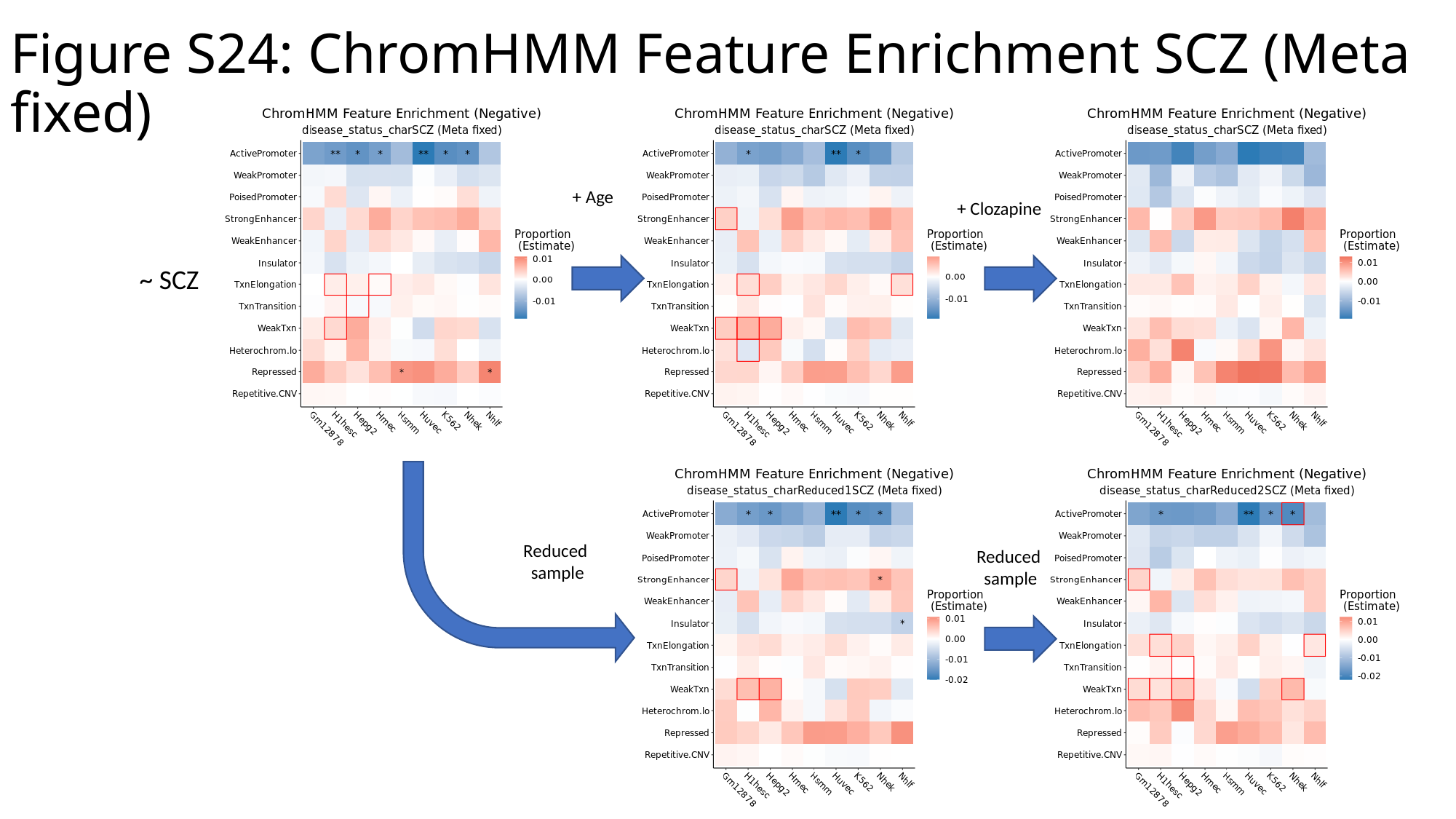

Figure S24: ChromHMM Feature Enrichment SCZ (Meta fixed)
+ Age
+ Clozapine
~ SCZ
Reduced
sample
Reduced
sample

### Slide 26
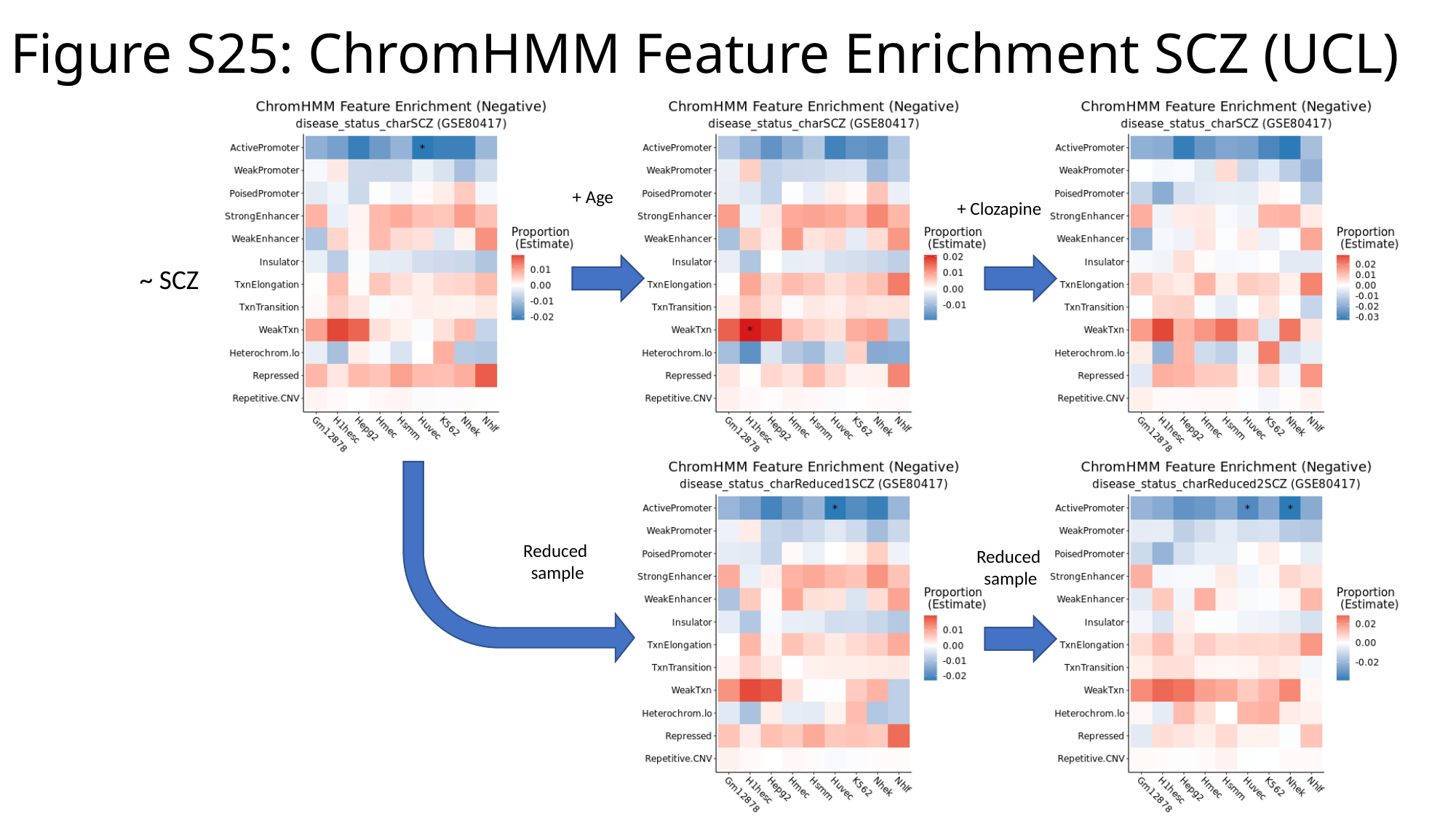

Figure S25: ChromHMM Feature Enrichment SCZ (UCL)
+ Age
+ Clozapine
~ SCZ
Reduced
sample
Reduced
sample

### Slide 27
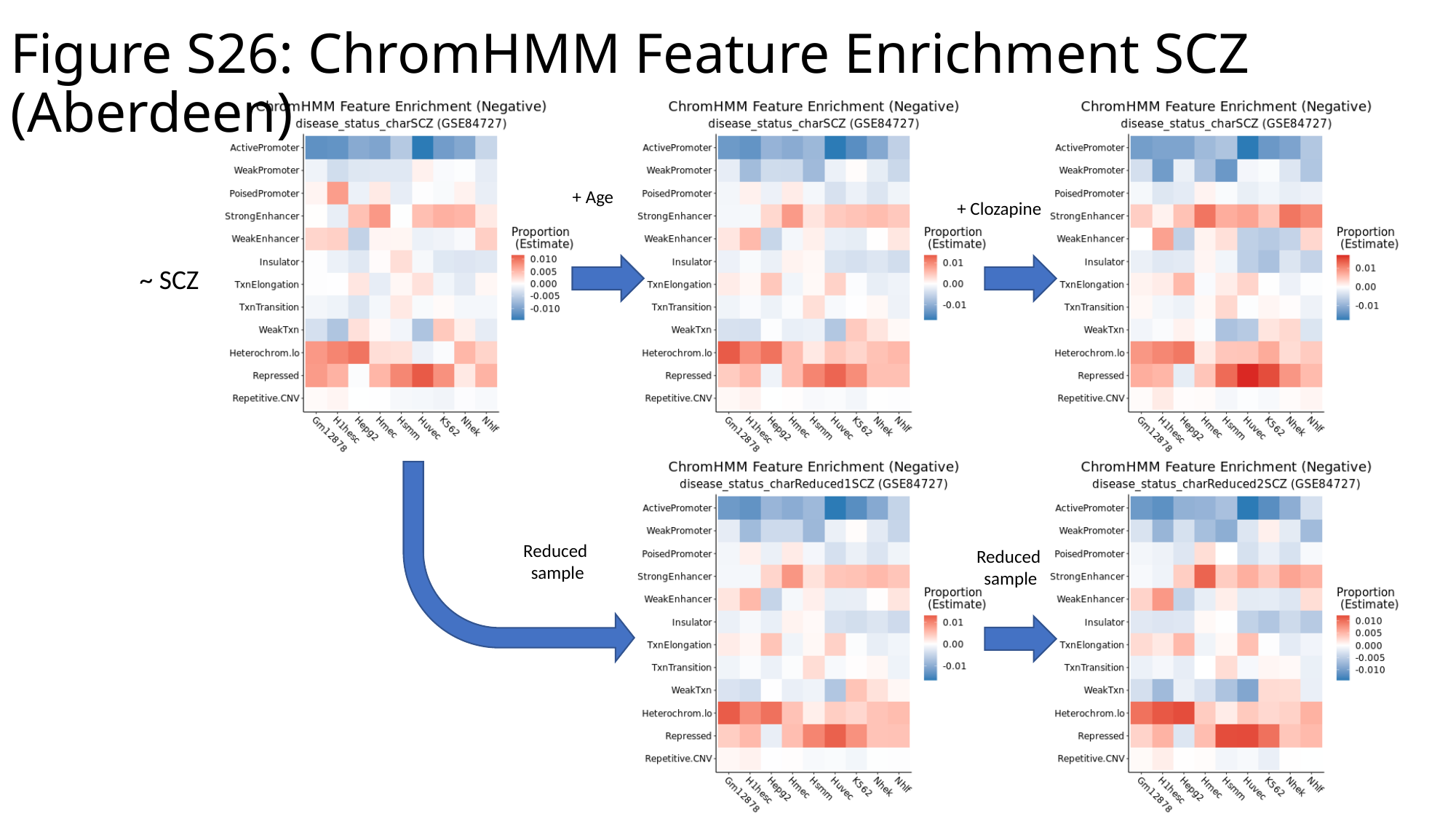

Figure S26: ChromHMM Feature Enrichment SCZ (Aberdeen)
+ Age
+ Clozapine
~ SCZ
Reduced
sample
Reduced
sample

### Slide 28
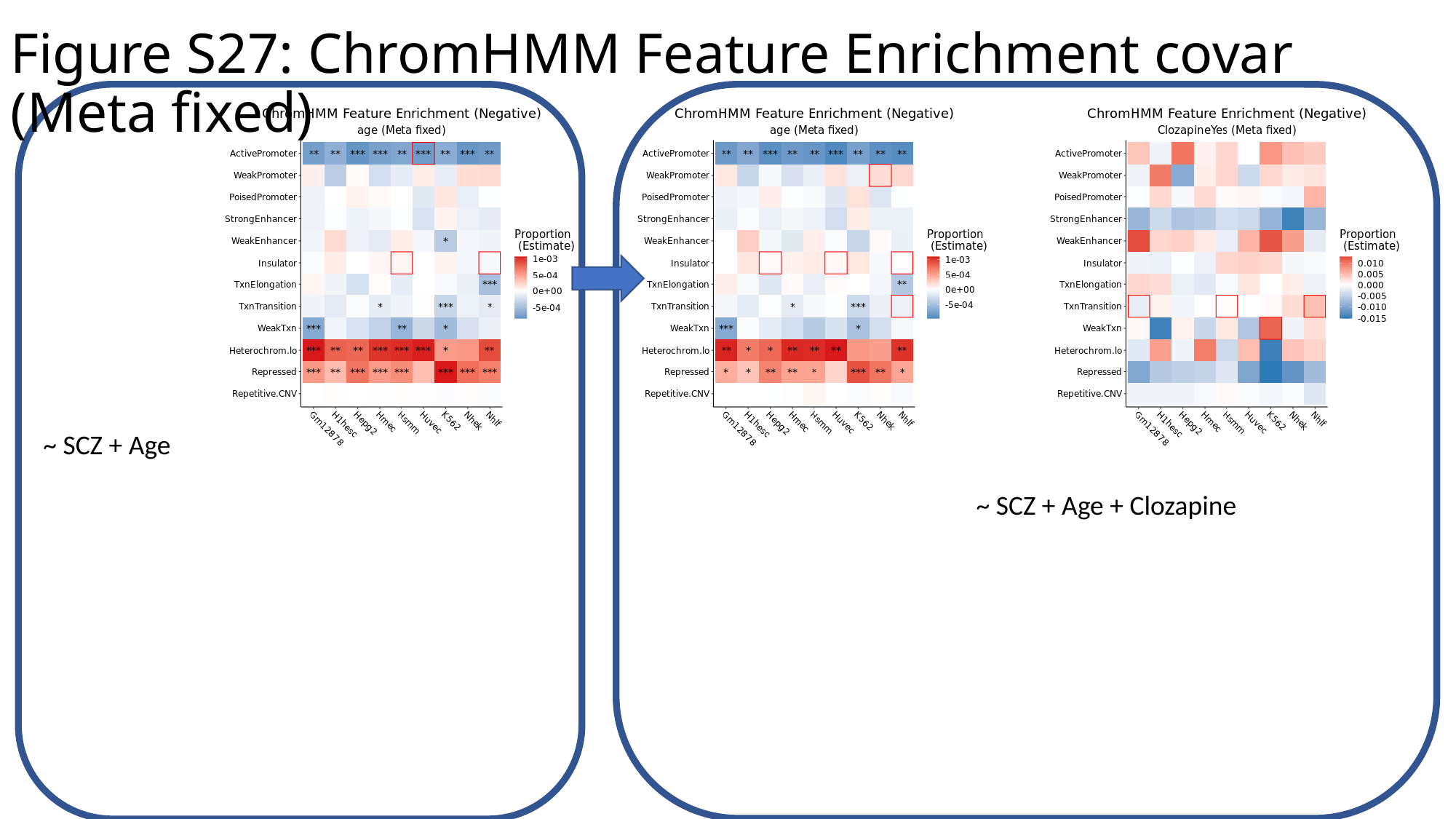

Figure S27: ChromHMM Feature Enrichment covar (Meta fixed)
~ SCZ + Age
~ SCZ + Age + Clozapine

### Slide 29
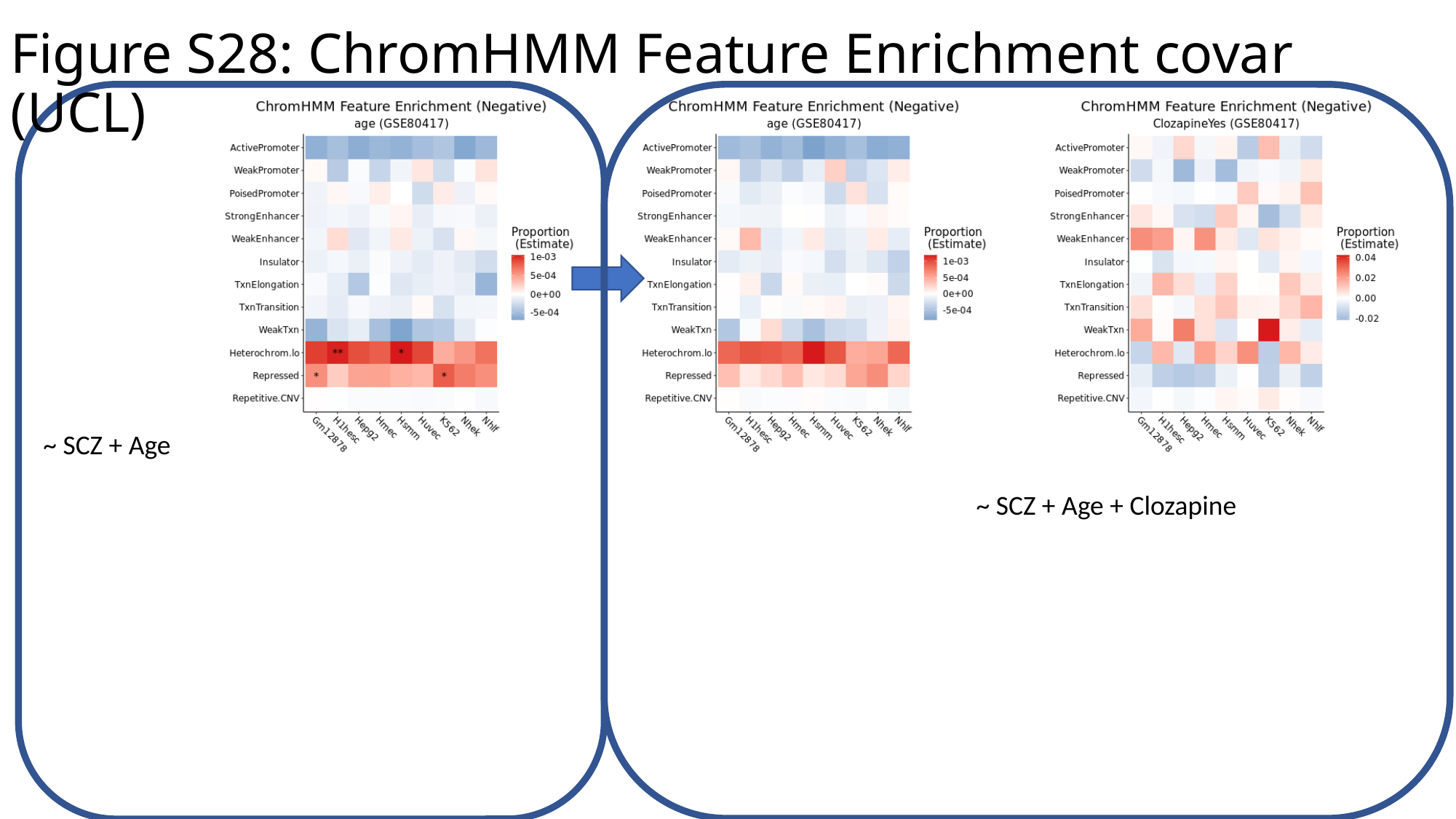

Figure S28: ChromHMM Feature Enrichment covar (UCL)
~ SCZ + Age
~ SCZ + Age + Clozapine

### Slide 30
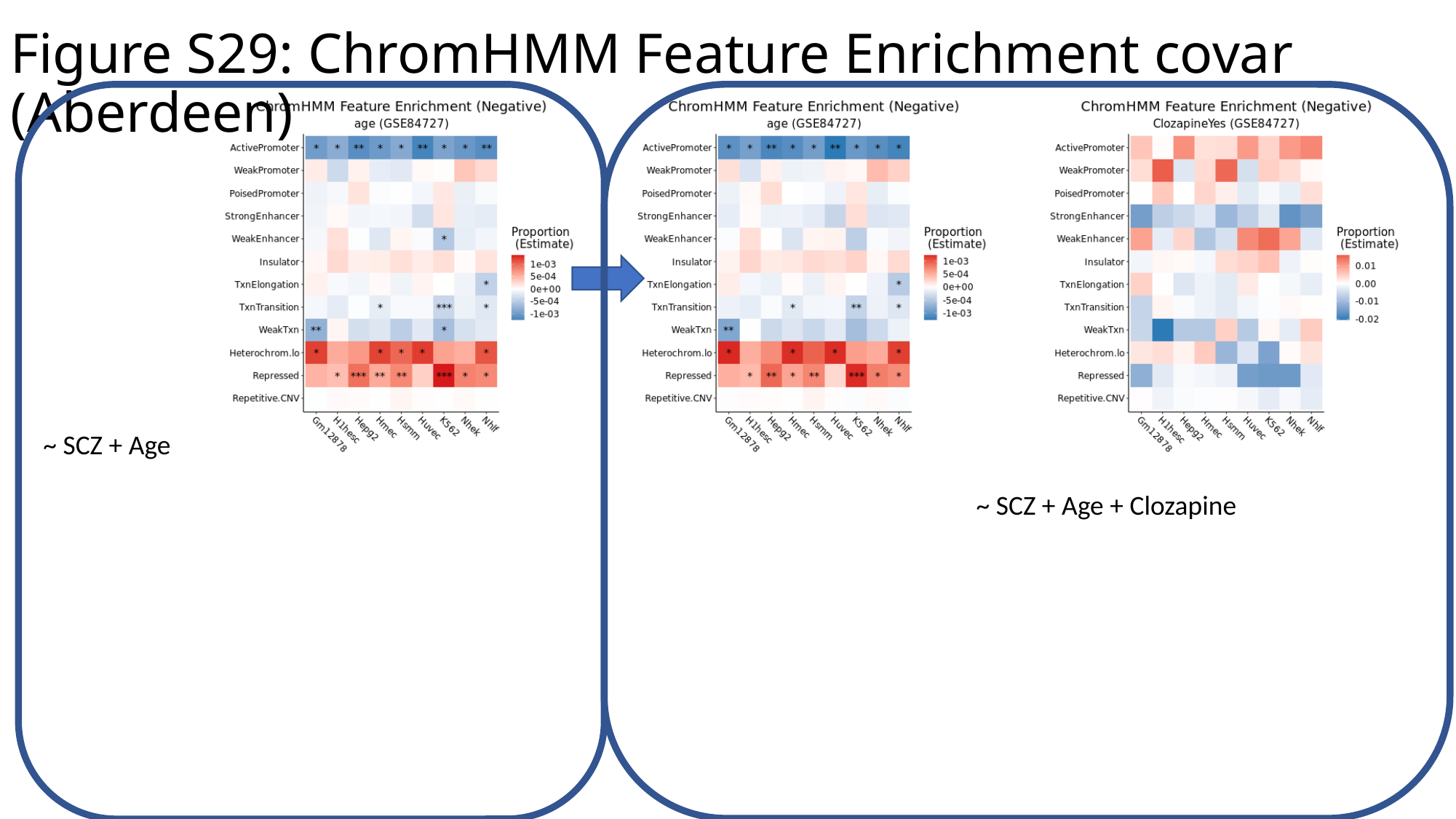

Figure S29: ChromHMM Feature Enrichment covar (Aberdeen)
~ SCZ + Age
~ SCZ + Age + Clozapine

### Slide 31
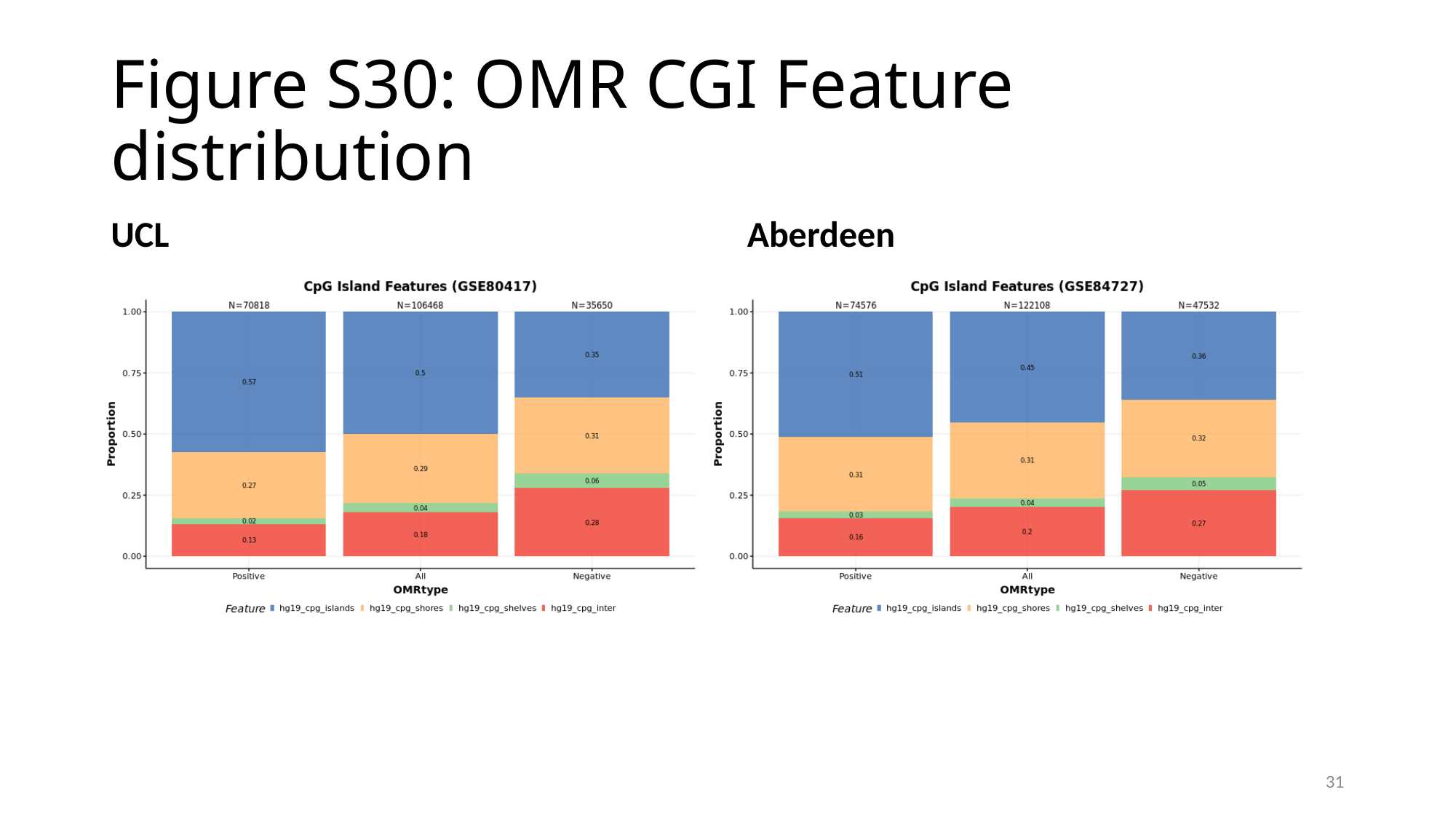

#### Figure S30: OMR CGI Feature distribution
UCL
Aberdeen
31

### Slide 32
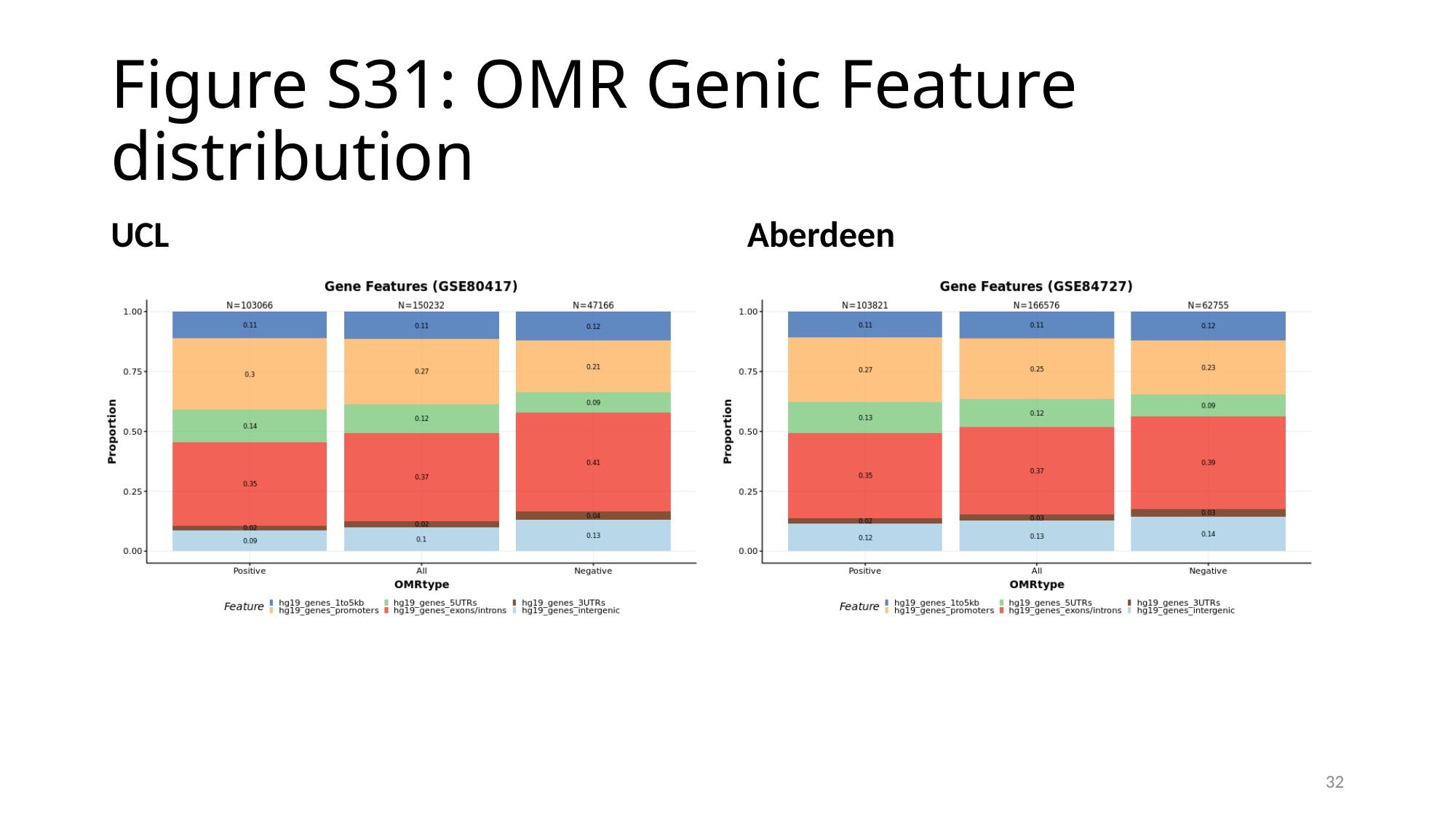

#### Figure S31: OMR Genic Feature distribution
UCL
Aberdeen
32

### Slide 33
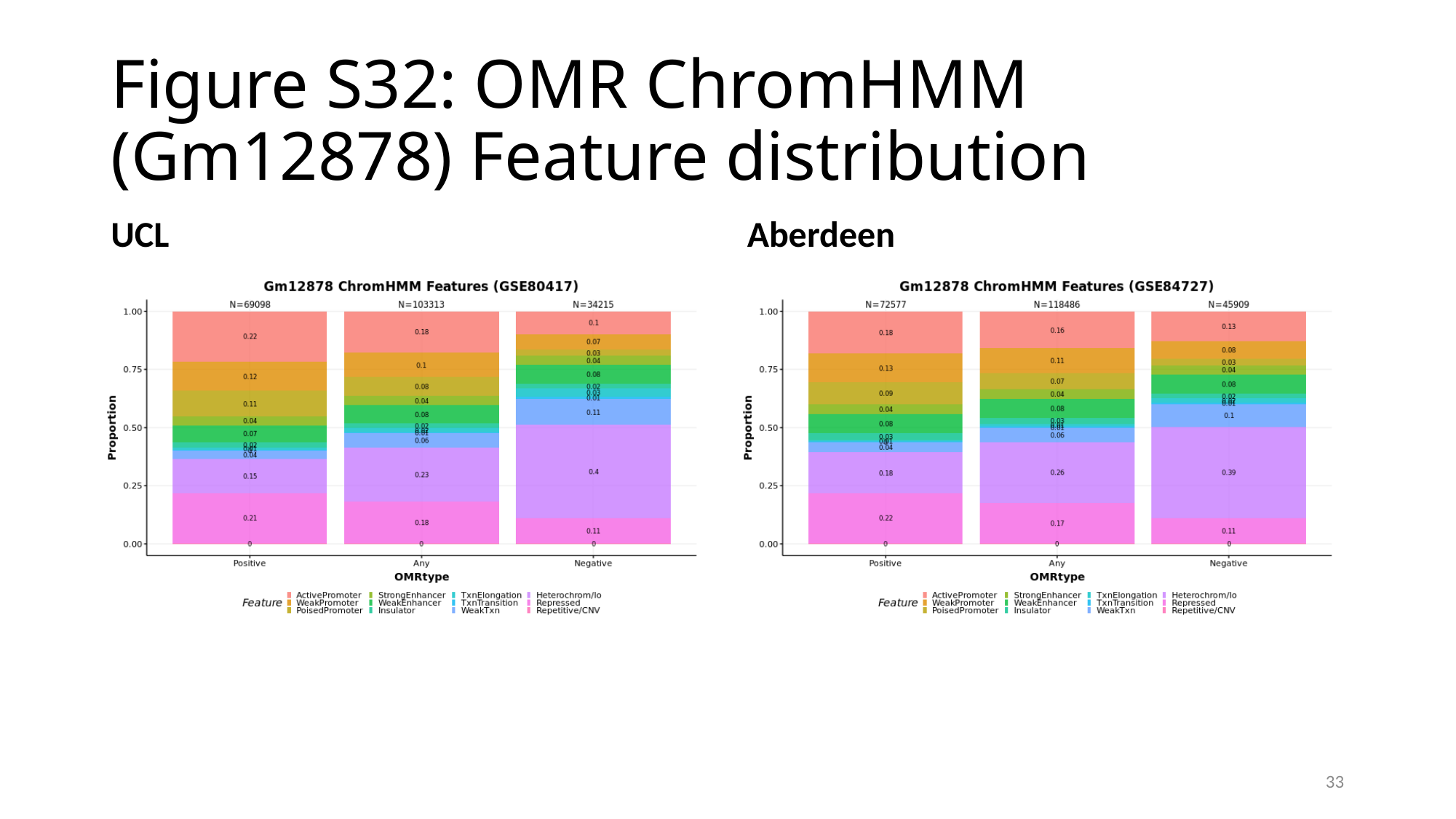

#### Figure S32: OMR ChromHMM (Gm12878) Feature distribution
UCL
Aberdeen
33

### Slide 34
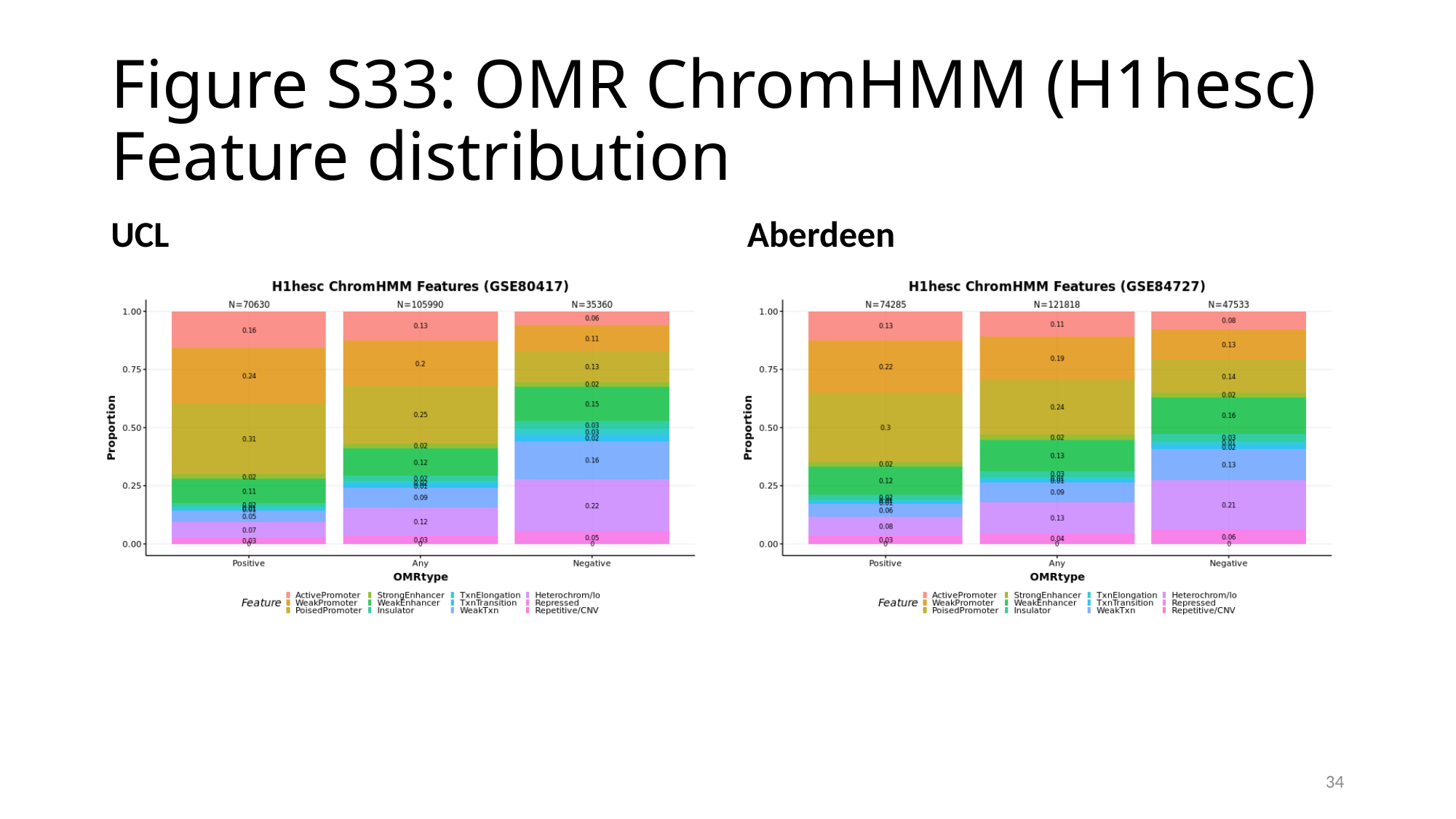

#### Figure S33: OMR ChromHMM (H1hesc) Feature distribution
UCL
Aberdeen
34

### Slide 35
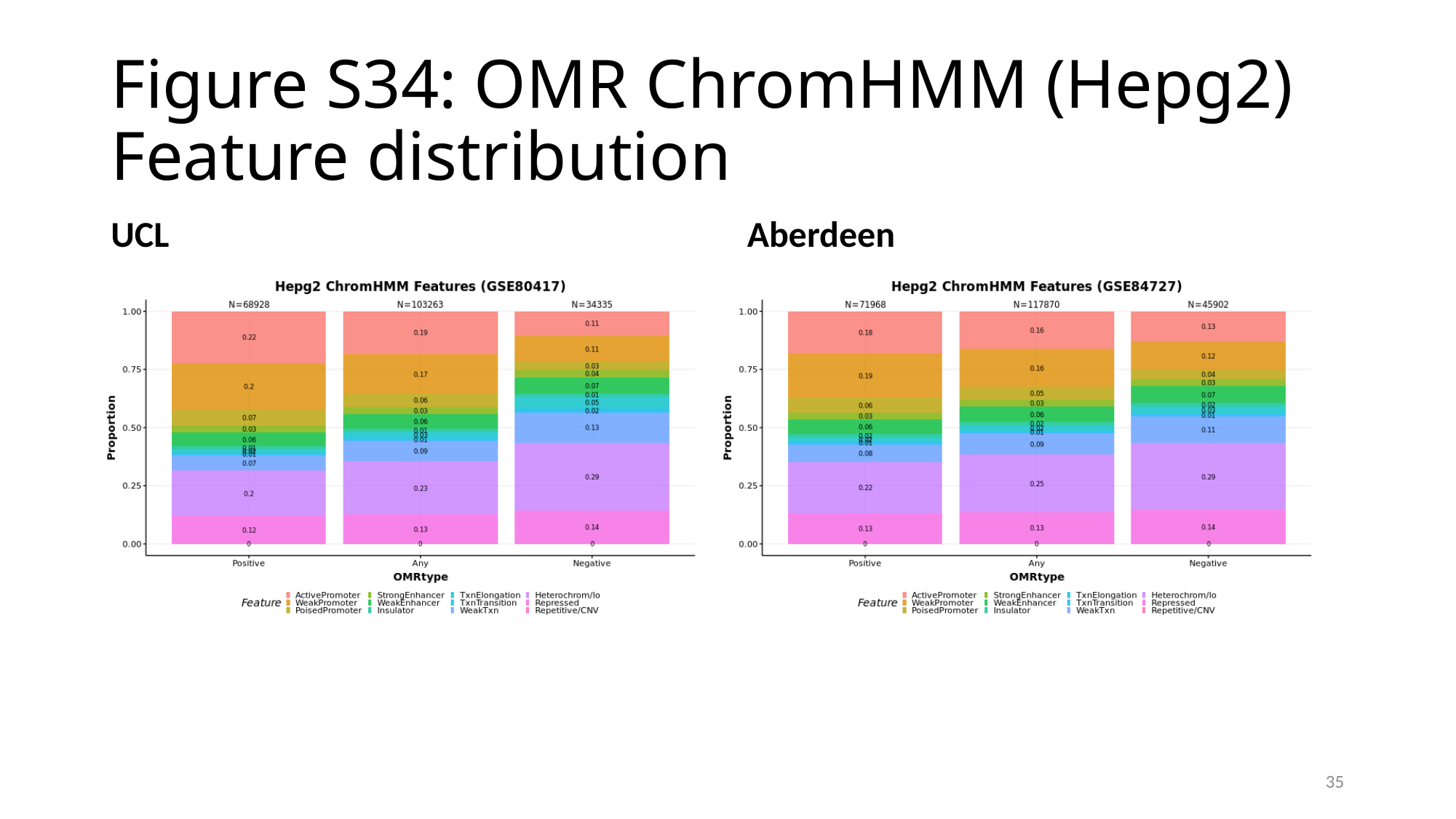

#### Figure S34: OMR ChromHMM (Hepg2) Feature distribution
UCL
Aberdeen
35

### Slide 36
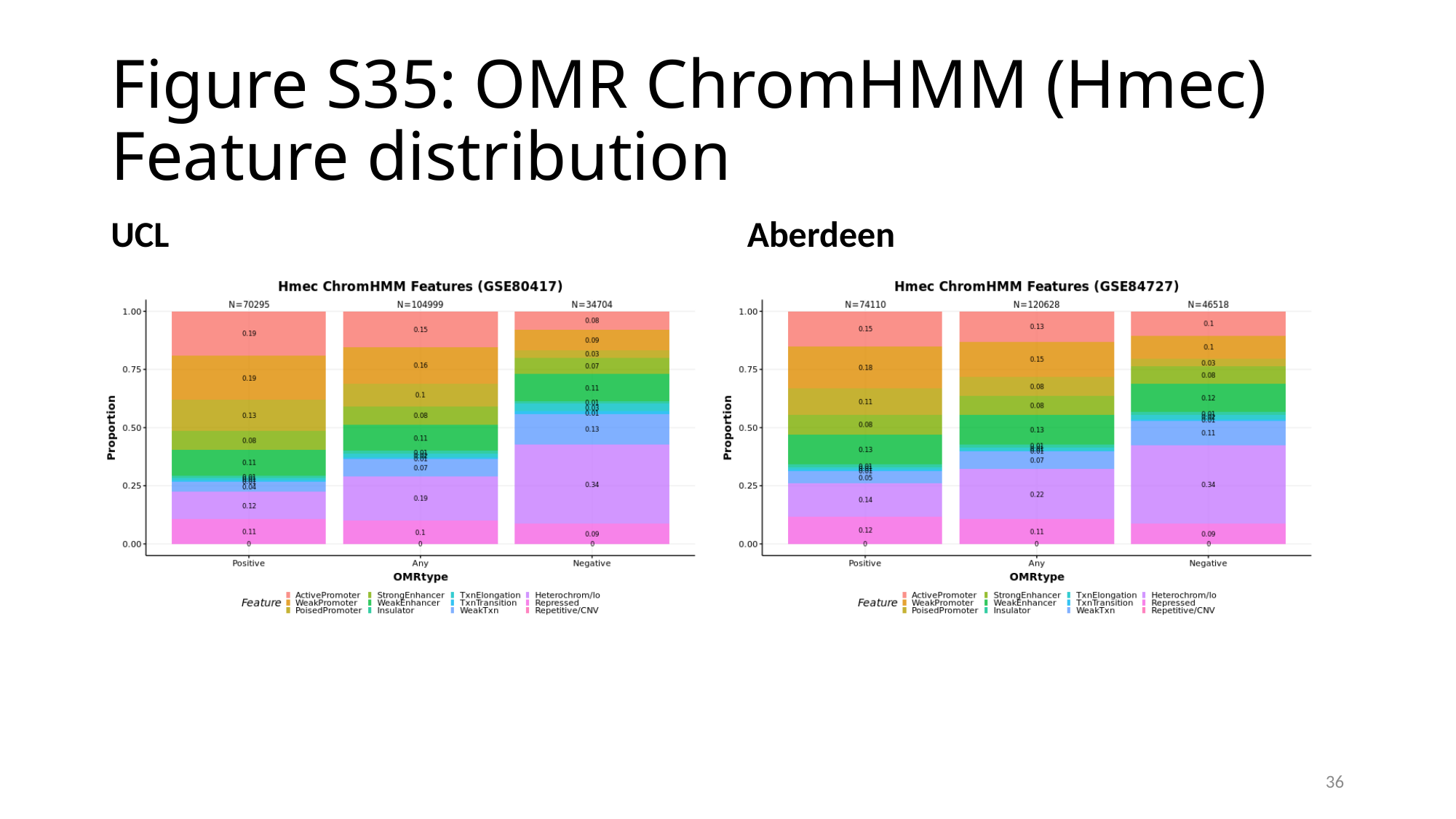

#### Figure S35: OMR ChromHMM (Hmec) Feature distribution
UCL
Aberdeen
36

### Slide 37
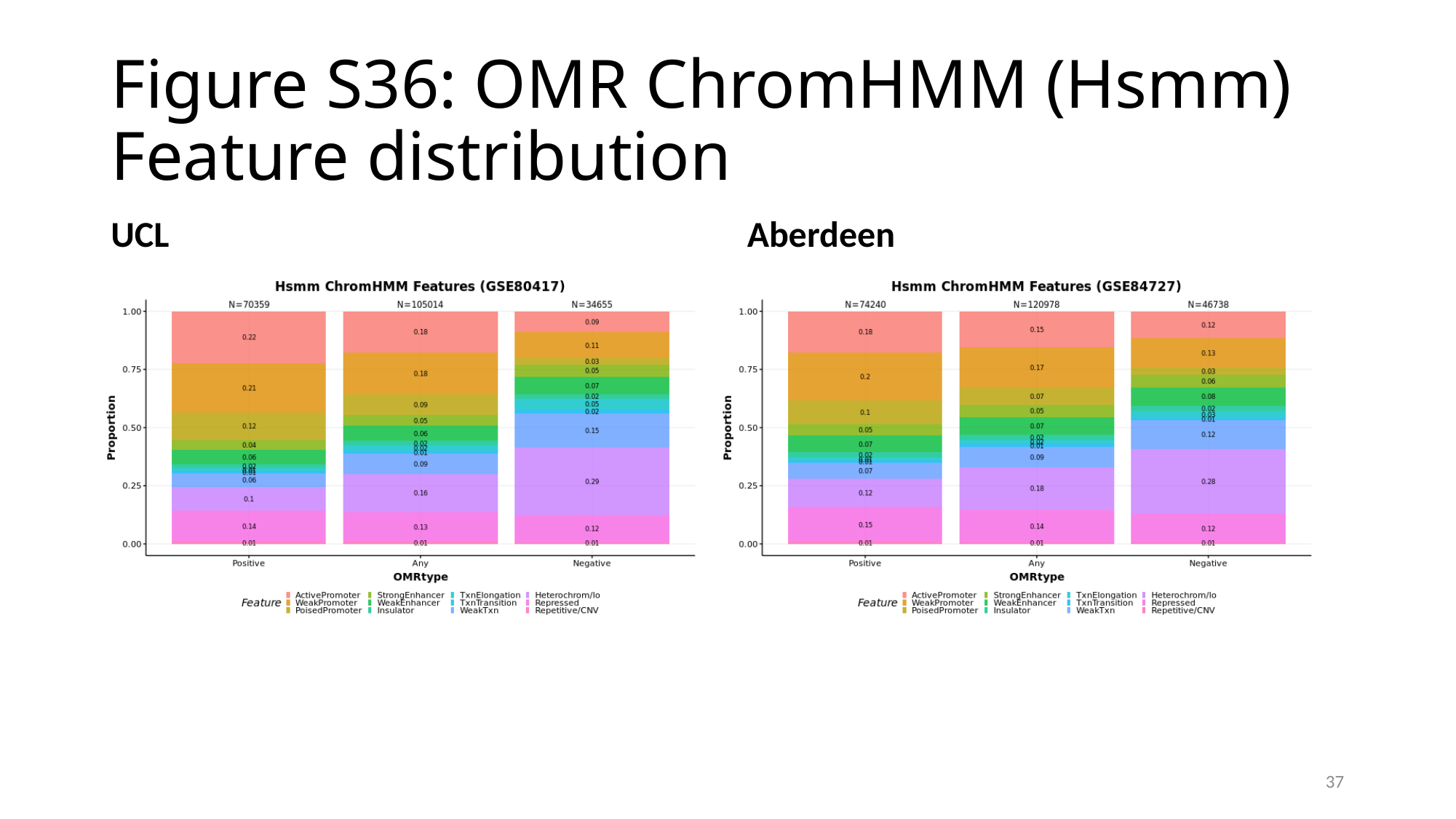

#### Figure S36: OMR ChromHMM (Hsmm) Feature distribution
UCL
Aberdeen
37

### Slide 38
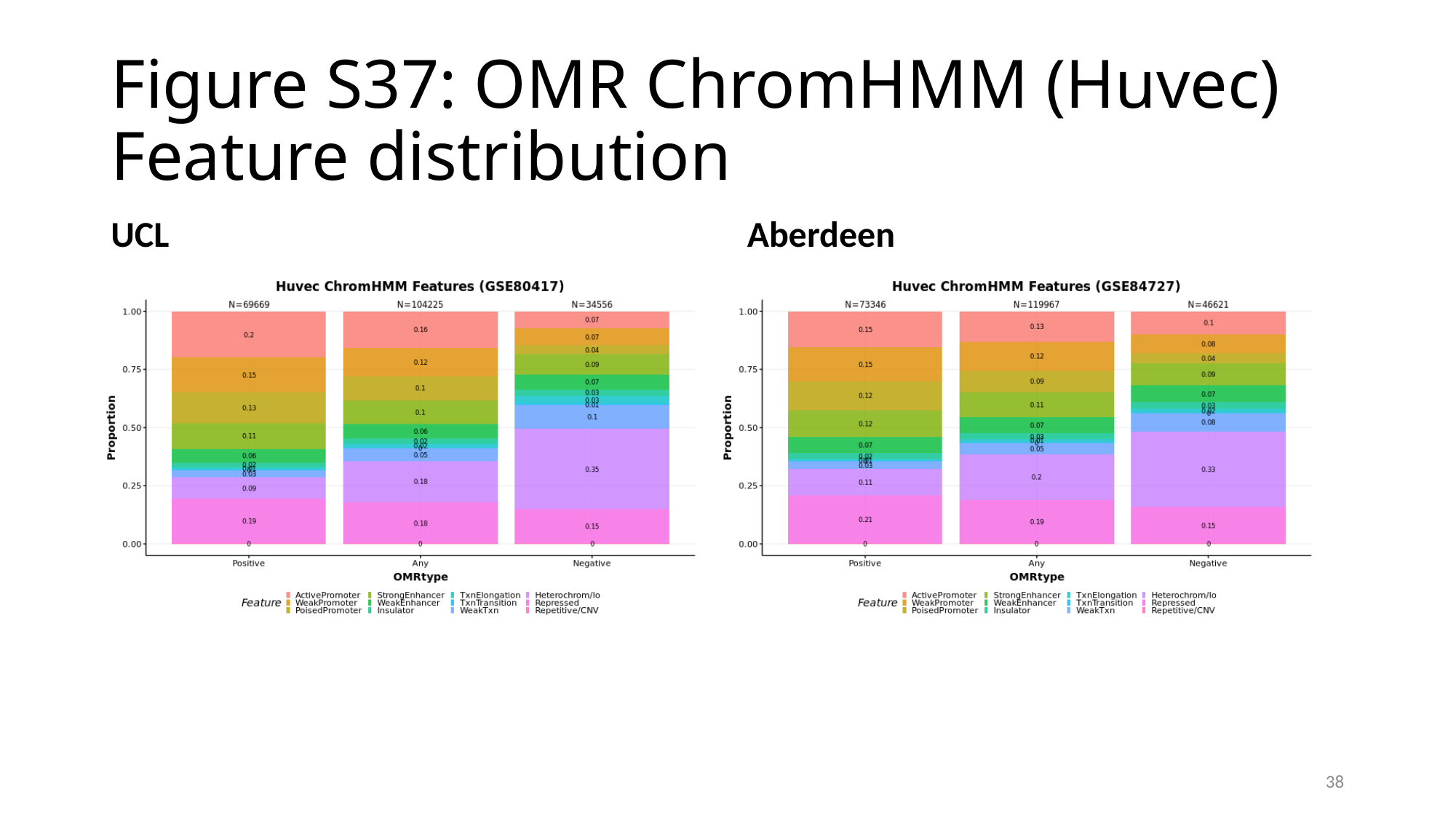

#### Figure S37: OMR ChromHMM (Huvec) Feature distribution
UCL
Aberdeen
38

### Slide 39
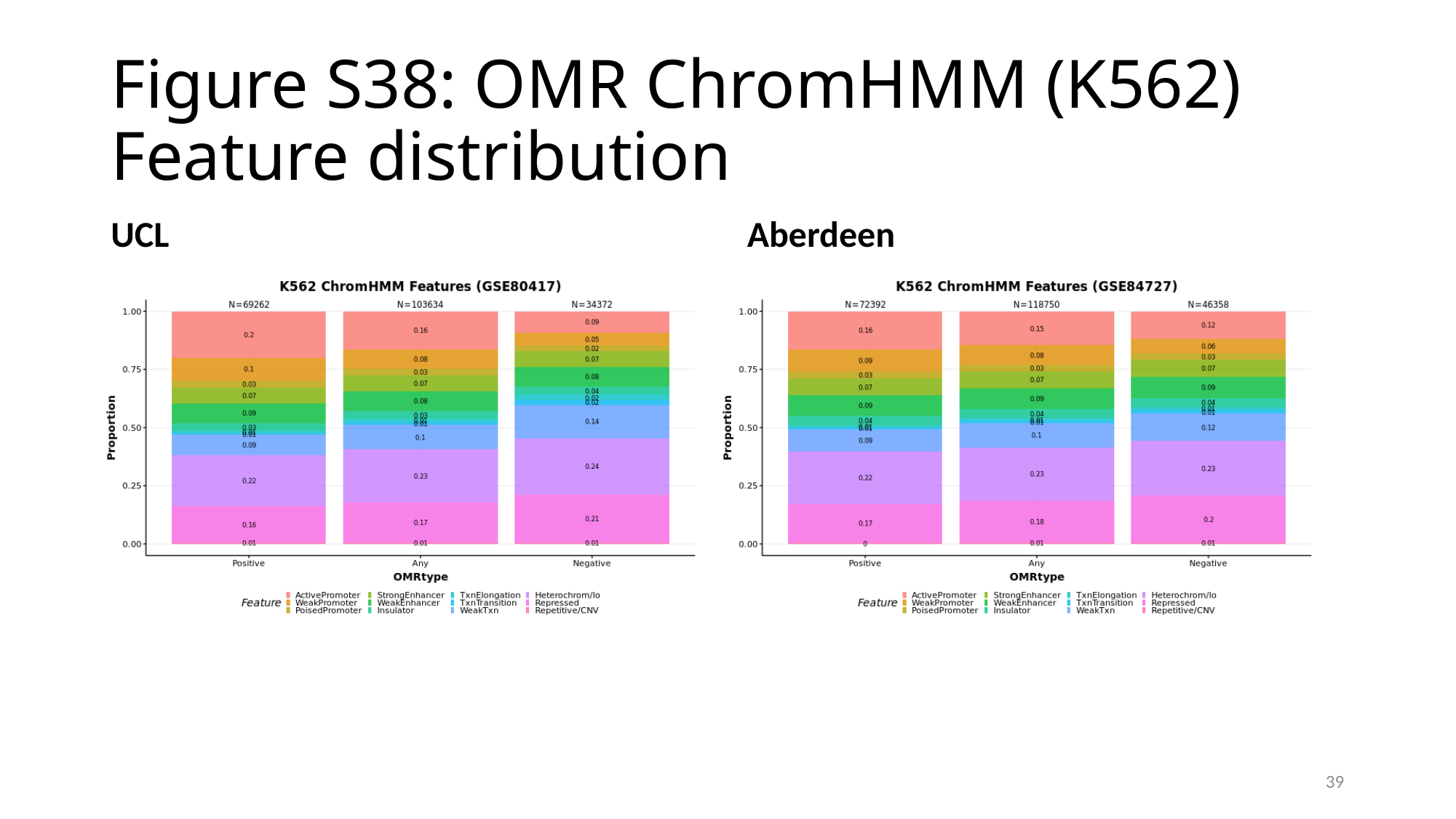

#### Figure S38: OMR ChromHMM (K562) Feature distribution
UCL
Aberdeen
39

### Slide 40
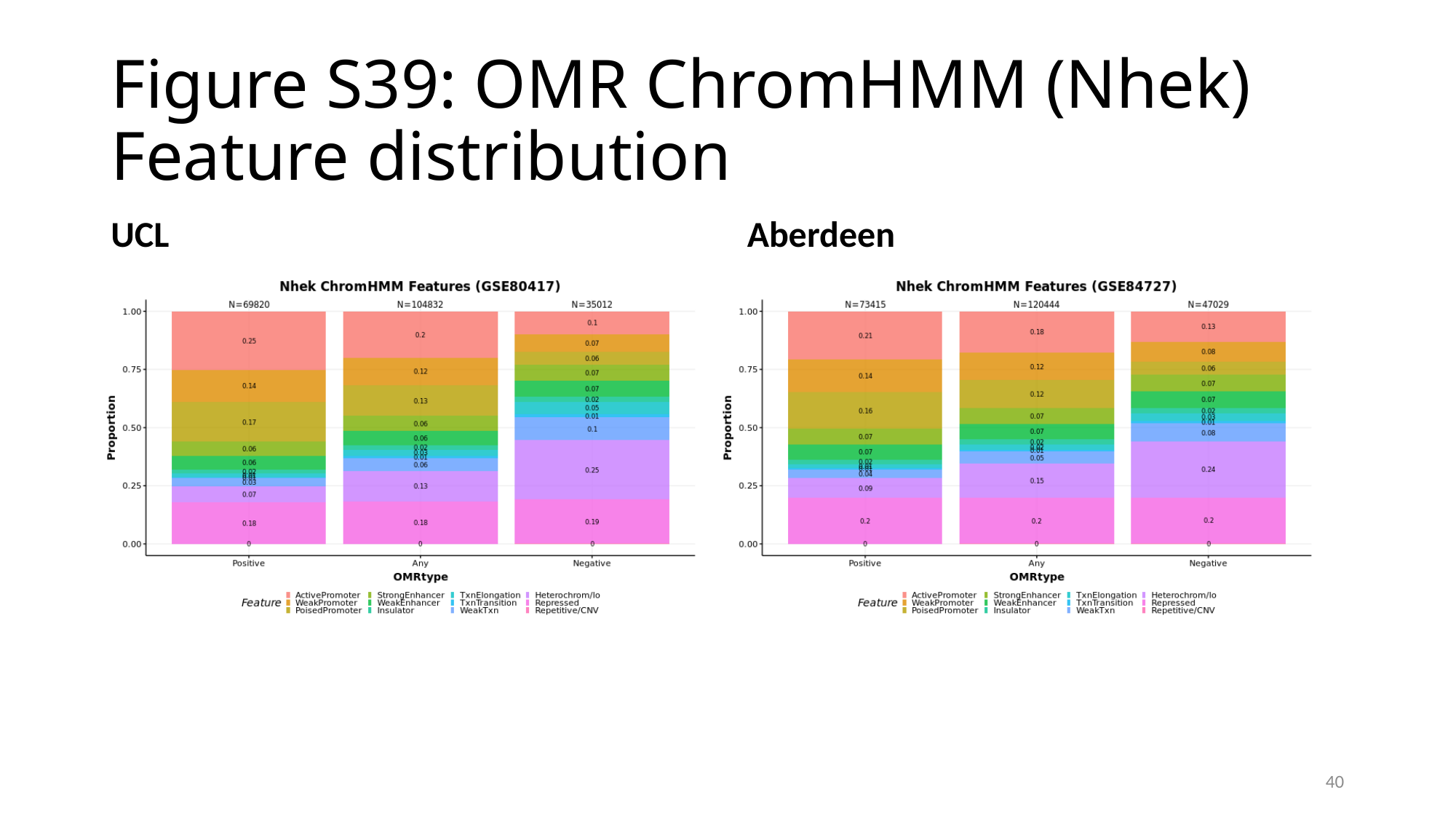

#### Figure S39: OMR ChromHMM (Nhek) Feature distribution
UCL
Aberdeen
40

### Slide 41
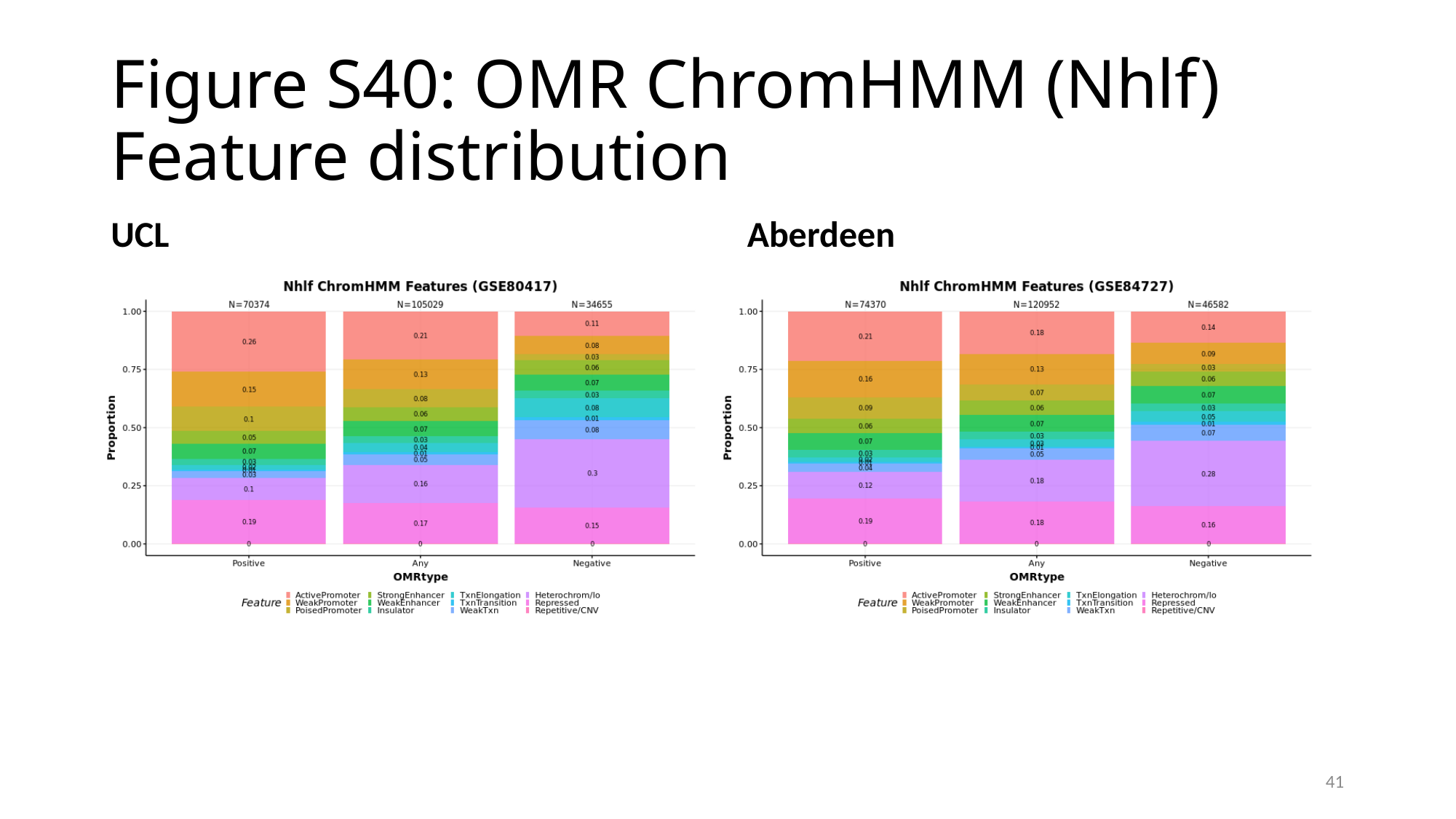

#### Figure S40: OMR ChromHMM (Nhlf) Feature distribution
UCL
Aberdeen
41
